# Cortical *Foxp2* supports behavioral flexibility and developmental dopamine D1 receptor expression

**DOI:** 10.1101/624973

**Authors:** Marissa Co, Stephanie L. Hickey, Ashwinikumar Kulkarni, Matthew Harper, Genevieve Konopka

## Abstract

Genetic studies have associated *FOXP2* variation with speech and language disorders and other neurodevelopmental disorders involving pathology of the cortex. In this brain region, *FoxP2* is expressed from development into adulthood, but little is known about its downstream molecular and behavioral functions. Here, we characterized cortex-specific *Foxp2* conditional knockout mice and found a major deficit in reversal learning, a form of behavioral flexibility. In contrast, they showed normal activity levels, anxiety, and vocalizations, save for a slight decrease in neonatal call loudness. These behavioral phenotypes were accompanied by decreased cortical dopamine D1 receptor (D1R) expression at neonatal and adult stages, while general cortical development remained unaffected. Finally, using single-cell transcriptomics, we identified at least five excitatory and three inhibitory D1R-expressing cell types in neonatal frontal cortex, and we found changes in D1R cell type composition and gene expression upon cortical *Foxp2* deletion. Strikingly, these alterations included non-cell-autonomous changes in upper-layer neurons and interneurons. Together these data support a role for *Foxp2* in the development of dopamine-modulated cortical circuits and behaviors relevant to neurodevelopmental disorders.

## Introduction

*FoxP2* encodes a forkhead box transcription factor required for proper brain development and function across species, particularly in neural circuits underlying vocalization and motor-skill learning (French and Fisher 2014; Konopka and Roberts 2016). In humans, *FOXP2* mutations cause a speech and language disorder characterized by childhood apraxia of speech and additional oral motor, linguistic, and/or cognitive deficits (Morgan et al. 2017; Schulze et al. 2017). Recent studies have broadened the clinical spectrum of *FOXP2* by identifying variants associated with autism spectrum disorder (ASD) and attention deficit/hyperactivity disorder (ADHD) (Demontis et al. 2019; Reuter et al. 2017; Satterstrom et al. 2019). Thus, *FOXP2* may subserve general neural functions impaired across neurodevelopmental disorders (NDDs).

*FoxP2* is expressed in the developing and mature cerebral cortex, a site of pathology in *FOXP2*-related speech and language disorders as well as in ASD (van Rooij et al. 2018; Vargha-Khadem et al. 2005). Here, *Foxp2* expression is specific to layer 6 corticothalamic projection neurons (CThPNs) and some layer 5 pyramidal tract neurons but excluded from intratelencephalic projection neurons (ITPNs) (Kast et al. 2019; Sorensen et al. 2015; Tasic et al. 2016). Acute manipulations of *Foxp2* expression in embryonic cortex have implicated this gene in cortical neurogenesis and neuronal migration (Garcia-Calero et al. 2016; Tsui et al. 2013). However, mice with cortical *Foxp2* deletion show overtly normal cortical histoarchitecture, suggesting that *Foxp2* may be dispensable for gross corticogenesis (French et al. 2018; Kast et al. 2019; Medvedeva et al. 2018). Nonetheless, these mice show abnormalities in social behavior and motor-skill learning, warranting further investigation into molecular and cellular processes disrupted by cortical *Foxp2* deletion (French et al. 2018; Medvedeva et al. 2018). Foxp2 has been shown to act upstream of two synaptic genes, *Srpx2* and *Mint2*, in cortical neurons, but little else is known about molecular networks regulated by Foxp2 specifically in the cortex (Medvedeva et al. 2018; Sia et al. 2013). Furthermore, it is unknown whether loss of cortical *Foxp2* causes additional NDD-relevant behavioral deficits, such as cognitive impairment or hyperactivity.

In this study, we characterized NDD-relevant behaviors and their potential underlying cellular and molecular mechanisms in cortex-specific *Foxp2* conditional knockout mice (*Emx1-Cre; Foxp2^flox/flox^*). We show that this deletion impaired reversal learning, a form of behavioral flexibility, while sparing other NDD-associated behaviors, such as vocal communication and hyperactivity. Using immunohistochemistry and genetic reporter mice, we confirmed grossly normal cortical development upon *Foxp2* deletion but found decreased expression of cortical dopamine D1 receptors at neonatal and adult stages. Last, using single-cell transcriptomics, we characterized neonatal dopamine D1 receptor-expressing neuronal subtypes, and we identified non-cell-autonomous effects of *Foxp2* deletion on interneuron development and upper-layer gene expression. Together these data support a role for *Foxp2* in specific aspects of cortical development potentially relevant to cognitive impairments seen in NDDs.

## Materials and Methods

### Mice

All procedures were approved by the Institutional Animal Care and Use Committee of UT Southwestern. *Emx1-Cre* (Gorski et al. 2002) (#005628, Jackson Laboratory), *Foxp2^flox/flox^* (French et al. 2007) (#026259, Jackson Laboratory) and *Drd1a-tdTomato* line 6 mice (Ade et al. 2011) (#016204, Jackson Laboratory, provided by Dr. Craig Powell) were backcrossed with C57BL/6J mice for at least 10 generations. Genotyping primers can be found in Supplemental Table 1. Experimental mice (*Foxp2* cKO) and control littermates were generated by crossing male *Emx1-Cre*; *Foxp2^flox/flox^* mice with female *Foxp2^flox/flox^* mice. When used, *Drd1a-tdTomato* was present in mice of either sex for breeding. Mice were group-housed under a 12 h light/dark cycle and given *ad libitum* access to food and water. Mice of both sexes were used for all experiments except adult USVs, which were measured in male mice.

**Table 1.**
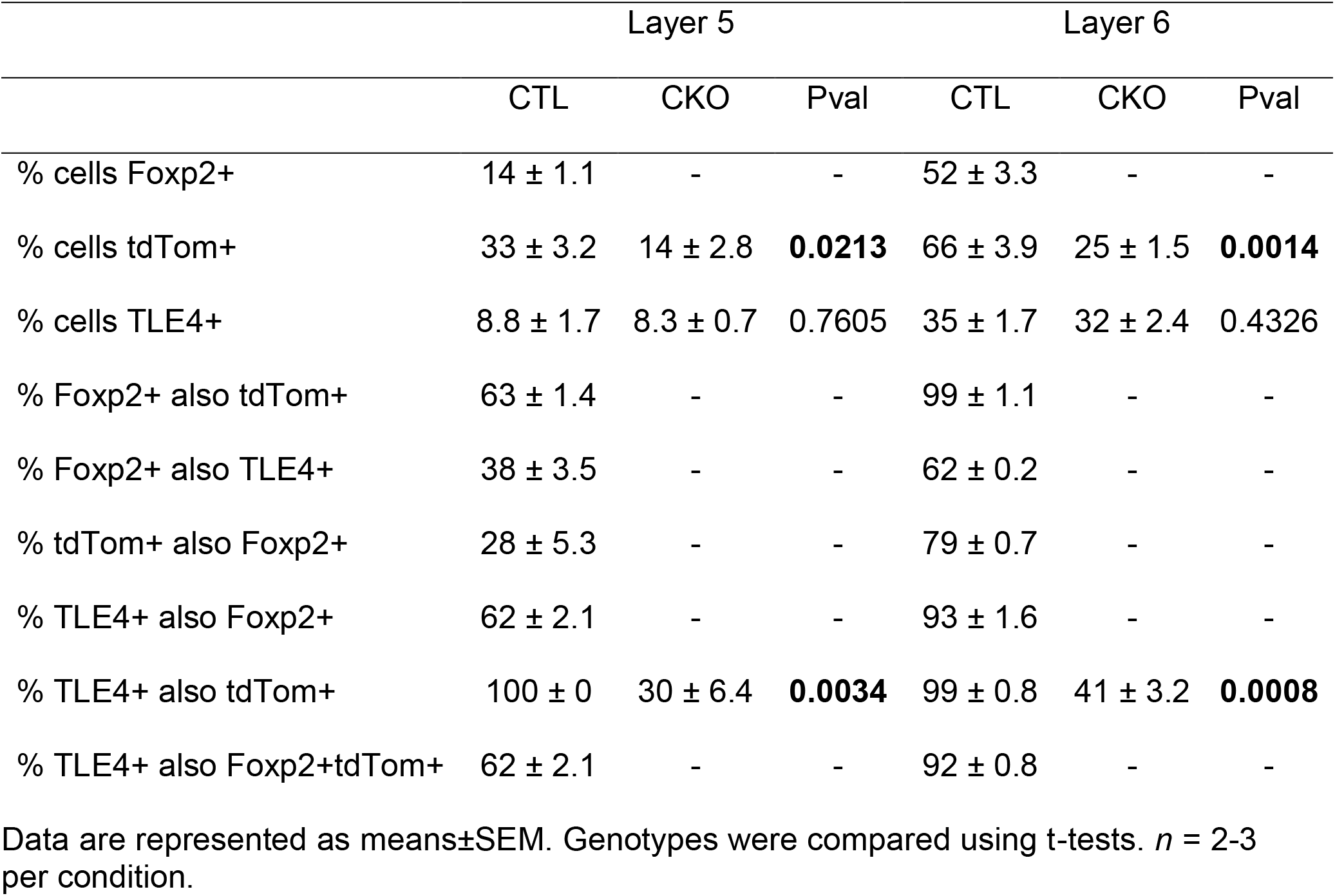
Summary of Foxp2, tdTomato, and TLE4 coexpression in P7 mPFC.

### Behavioral analyses

Adult *Foxp2* cKO and control littermates were tested at age 10-20 weeks, and pups were tested at postnatal days 4, 7, 10, and 14. Additional behavioral procedures can be found in the Supplemental Material.

### Reversal learning in water Y-maze

Mice were tested according to (Stoodley et al. 2017) using a Y-shaped apparatus filled with 20-22°C water up to 0.5 cm above a clear movable platform. On day 1, mice were habituated to the maze for 1 min without the platform. On days 2-4 (Training 1-3), mice were given 15 trials/day, up to 30 s each, to learn the platform location in one of the maze arms. Platform location was counter-balanced by cage to control for side biases. On days 4-6 (Reversal 1-3), the platform was moved to the opposite arm and mice were given 15 trials/day to learn the new location. The fraction of correct trials per day was calculated, as well as number of trials to reach a criterion of 5 consecutive correct trials. Differences between genotypes were assessed using a two-way ANOVA with Bonferroni’s multiple comparisons test.

### Adult courtship ultrasonic vocalizations

Mice were tested according to (Araujo et al. 2017). Male test mice were paired with age-matched C57BL/6J females for 1 week, then single-housed for 1 week. On the test day, males were habituated in their home cages to the testing room for 30 min, during which their cage lids were replaced with Styrofoam lids containing UltraSoundGate condenser microphones (Avisoft Bioacoustics) positioned at a fixed height of 20 cm. The microphones were connected to UltraSoundGate 416H hardware (Avisoft Bioacoustics) connected to a computer running RECORDER software (Avisoft Bioacoustics). At the start of testing, an unmated age-matched C57BL/6J female was placed in each cage and resultant male songs were recorded for 3 min. Spectrogram preparation and call detection were performed using MATLAB code developed by (Rieger and Dougherty 2016) based on methods from (Holy and Guo 2005). Differences between genotypes were assessed using unpaired t-tests. Call repertoire analysis was performed using the MUPET MATLAB package (Van Segbroeck et al. 2017) with a repertoire size of 100 units.

### Neonatal isolation ultrasonic vocalizations

Mice were tested according to (Araujo et al. 2015). After habituation in their home cages to the testing room for 30 min, individual pups were placed in plastic containers within 1 of 4 soundproof Styrofoam boxes with lids containing UltraSoundGate condenser microphones. Pups were randomly assigned to recording boxes at each postnatal time point. Isolation USVs were recorded for 3 min and analyzed using the same MATLAB code used for adult USV analysis (Rieger and Dougherty 2016). Other than call number, USV features were only computed for pups emitting at least 10 calls during the recording session. Differences between genotypes were assessed using a two-way ANOVA with Bonferroni’s multiple comparisons test.

### Immunohistochemistry

Neonatal and adult mice were anesthetized (pups by cryoanesthetization, adults by injection with 80-100 mg/kg Euthasol) and transcardially perfused with 4% PFA, and their brains were post-fixed overnight in 4% PFA. After cryoprotection in 30% sucrose overnight, brains were embedded in Tissue-Tek CRYO-OCT Compound (#14-373-65, Thermo Fisher Scientific) and cryosectioned at 20-40 μm. Staining was performed on free-floating sections and all washes were performed with TBS or 0.4% Triton X-100 in TBS (TBS-T) unless otherwise stated. For TLE4 staining, antigen retrieval was performed in citrate buffer (10 mM tri-sodium citrate, 0.05% Tween-20, pH 6) for 10 min at 95°C. Free aldehydes were quenched with 0.3M glycine in TBS for 1 h at room temperature. Sections were incubated overnight at 4°C in primary antibodies diluted in 3% normal donkey serum and 10% bovine serum albumin (BSA) in TBS-T. Secondary antibody incubations were performed for 1 h at room temperature in 10% BSA in TBS-T. Sections were mounted onto slides, incubated in DAPI solution (600 nM in PBS) for 5 min at room temperature, and washed 3X with PBS. Coverslips were mounted using ProLong Diamond Antifade Mountant (#P36970, Thermo Fisher Scientific). The following antibodies and dilutions were used: mouse α-β-actin (#A1978, Millipore Sigma, 10 μg/ml), rabbit α-β-tubulin (#ab6046, Abcam, 1:500), mouse α-DARPP-32 (#sc-271111, Santa Cruz Biotechnology, 1:250), rabbit α-FOXP2 (#5337S, Cell Signaling Technology, 1:250), rabbit a-SP9 (#PA564038, Thermo Fisher Scientific, 1:100), goat α-tdTomato (#LS-C340696, LifeSpan BioSciences, 1:500), mouse α-TLE4 (#sc-365406, Santa Cruz Biotechnology, 1:200), species-specific secondary antibodies produced in donkey and conjugated to Alexa Fluor 488, Alexa Fluor 555, or Alexa Fluor 647 (Thermo Fisher Scientific, 1:2000).

### Imaging and Image Analysis

Images were acquired using a Zeiss LSM 880 confocal microscope at the UT Southwestern Neuroscience Microscopy Facility and processed and analyzed using Zeiss ZEN Lite and FIJI. For quantifications, tile scan Z-stack images of the region of interest were acquired at 20X magnification from similar coronal sections across 2-3 mice/genotype. Stitched maximum intensity projection images were used for manual cell counting using the FIJI Cell Counter plugin or for fluorescence intensity measurements using the FIJI Plot Profile function. For tdTomato+ cell counts by layer, layers in mPFC were defined based a combination of DAPI-based cytoarchitecture and TLE4+ cell distribution. Differences between genotypes were assessed using a two-way ANOVA with Bonferroni’s multiple comparisons test.

### Western blotting

Western blotting was performed as previously described (Araujo et al. 2015). Frontal cortex tissue from 3 mice/genotype at P7 was lysed in RIPA buffer containing protease inhibitors. Protein concentrations were determined via Bradford assay (Bio-Rad Laboratories) and 50 μg protein per sample were run on an SDS-PAGE gel and transferred to an Immun-Blot PVDF Membrane (Bio-Rad Laboratories) using standard protocols. The following antibodies and dilutions were used: rabbit α-DARPP-32 (#AB10518, Millipore Sigma, 1 μg/ml), mouse α-GAPDH (#MAB374, Millipore Sigma, 1:10,000), donkey α-rabbit IgG IRDye 800 (#926-32213, LI-COR Biosciences, 1:20,000), donkey α-mouse IgG IRDye 680 (#926-68072, LI-COR Biosciences, 1:20,000). Blots were imaged using an Odyssey Infrared Imaging System (LI-COR Biosciences).

## Single-cell RNA-seq (scRNA-seq)

### Tissue processing and library generation

Tissue was dissociated for scRNA-seq based on (Tasic et al. 2016). P7 mice were sacrificed by rapid decapitation and brains were quickly removed and placed in ice-cold artificial cerebrospinal fluid (ACSF) (126 mM NaCl, 20 mM NaHCO_3_, 20 mM D-Glucose, 3 mM KCl, 1.25 mM NaH_2_PO_4_, 2 mM CaCl_2_, 2 mM MgCl_2_) bubbled with 95% O_2_ and 5% CO_2_. 400-μm coronal sections were made in ACSF using a VF-200 Compresstome and transferred to a room temperature recovery chamber with ACSF containing channel blockers DL-AP5 sodium salt (50 μM), DNQX (20 μM), and tetrodotoxin (100 nM) (ACSF+). After 5 min, frontal isocortex was separated from olfactory areas, cut into smaller pieces and incubated in 1 mg/ml pronase (#P6911, Sigma-Aldrich) in ACSF+ for 5 min. Pronase solution was replaced with 1% BSA in ACSF and tissue pieces were gently triturated into single-cell suspension using polished glass Pasteur pipettes with 600 μm, 300 μm, and 150 μm openings. Cells were filtered twice through Flowmi 40 μm Cell Strainers (#H13680-0040, Bel-Art) and live, single tdTomato+ cells were sorted using a BD FACSAria (BD Biosciences) at the UT Southwestern Flow Cytometry Facility. After sorting, cells were centrifuged and resuspended in 0.04% BSA in ACSF to target 1000 cells/sample using the Chromium Single Cell 3’ Library & Gel Bead Kit v2 (#120237, 10x Genomics) (Zheng et al. 2017). Tissue and library preparation were performed in the following batches: Batch 1 – D1Tom-CTL1 (F), D1Tom-CKO1 (F); Batch 2 – D1Tom-CTL2 (F), D1Tom-CKO2 (M). Libraries were sequenced using an Illumina NextSeq 500 at the McDermott Sequencing Core at UT Southwestern.

### Data processing

BCL files were demultiplexed with the i7 index using Illumina’s bcl2fastq v2.17.1.14 and *mkfastq* from 10x Genomics CellRanger v2.1.1. Extracted paired-end fastq files, consisting of a 26 bp cell barcode and unique molecular identifier (UMI) (R1) and a 124 bp transcript sequence (R2), were checked for read quality using FASTQC v0.11.5 (Andrews 2010). R1 reads were used to estimate and identify real cells using *whitelist* from UMI-tools v0.5.4 (Smith et al. 2017). A whitelist of cell barcodes and R2 fastq files were used to extract reads corresponding to cells using *extract* from UMI-tools v0.5.4. This step also appended the cell barcode and UMI sequence information from R1 to read names in the R2 fastq file. Extracted R2 reads were aligned to the mouse reference genome (MM10/GRCm38p6) from the UCSC genome browser (Kent et al. 2002) and reference annotation (Gencode vM17) using STAR v2.5.2b (Dobin et al. 2013) allowing up to 5 mismatches. Uniquely mapped reads were assigned to exons using *featureCounts* from the Subread package (v1.6.2) (Liao et al. 2014). Assigned reads were sorted and indexed using Samtools v1.6 (Liao et al. 2014) and then used to generate raw expression UMI count tables using *count* from UMI-tools v0.5.4. For libraries sequenced in multiple runs, the commonly identified cell barcodes between runs were used for downstream analysis.

### Clustering analysis

Cell clusters were identified using the Seurat R package (Butler et al. 2018). Individual cells were retained in the dataset based on the following criteria: <20,000 UMIs, <10% mitochondrial transcripts, <20% ribosomal protein gene transcripts. Sex chromosome and mitochondrial genes were removed from the analysis after filtering. The filtered data were log normalized with a scale factor of 10,000 using *NormalizeData*, and 1576 variable genes were identified with *FindVariableGenes* using the following parameters: mean.function = ExpMean, dispersion.function = LogVMR, x.low.cutoff = 0.2, x.high.cutoff = 2.5, y.cutoff = 0.5. Cell cycle scores were calculated using *CellCycleScoring* as per the Satija Lab cell cycle vignette (https://satijalab.org/seurat/cell_cycle_vignette.html). UMI number, percent mitochondrial transcripts, percent ribosomal protein gene transcripts, library, and cell cycle scores were regressed during scaling. Using JackStraw analysis, we selected principal components (PCs) 1-47 for clustering, excluding PCs with >1 immediate early gene (IEG) in the top 30 associated genes. We used a resolution of 1.6 for UMAP clustering and *ValidateClusters* with default parameters did not lead to cluster merging.

### Cell type annotation

Cluster marker genes were identified using *FindAllMarkers* with default parameters. Clusters were broadly annotated by enriched expression of canonical marker genes (e.g. Astrocytes: *Aqp4*; Microglia: *P2ry12*; Neurons: *Rbfox1*; Excitatory neurons: *Slc17a7*; Layer 2-4 neurons: *Satb2*; L5-6 neurons: *Fezf2, Tbr1*; L1 neurons: *Lhx5*; Interneurons: *Gad1*; Oligodendrocytes: *Sox10*). We refined these annotations by comparing our cluster markers with markers from a published scRNA-seq dataset from P0 mouse cortex (Loo et al. 2019). Metadata and raw expression values for this dataset were downloaded from https://github.com/jeremymsimon/MouseCortex. Cells were filtered as in the original publication and expression values normalized using Seurat’s *NormalizeData* with default parameters. The cluster identity of each cell was imported from the published metadata and cluster marker genes were identified using *FindAllMarkers* in Seurat. Enrichment of significant P0 marker genes (adj p<0.05) among our P7 cluster marker genes was analyzed using hypergeometric testing with a background of 2800 genes (the average of the median number of expressed genes in each cluster). P values were corrected for multiple comparisons using the Benjamini-Hochberg procedure.

### Neuronal re-clustering and annotation

Cells belonging to neuronal clusters (Clusters 2, 6, 8, 10-12, 14-17 in Supplemental Fig. 4B) were pulled from the full dataset and re-clustered with resolution 1.2 and PCs 1-59, excluding PCs with >1 IEG in the top 30 associated genes. Cell type annotation was performed as described above and refined using scRNA-seq marker genes identified in adult anterior lateral motor cortex (Tasic et al. 2018) (http://celltypes.brain-map.org/rnaseq/mouse). Two neuronal clusters (Clusters 2, 7) with enrichment of glial, mitochondrial, and/or ribosomal genes among their marker genes were included in analyses but excluded from data visualizations. Contributions of neurons to each cluster by genotype were compared using Fisher’s exact test.

### Differential gene expression analyses

We used the Wilcoxon rank sum test to calculate pseudo-bulk RNA-seq differentially expressed genes (DEGs) in two approaches: between genotypes for all neurons or between genotypes within each neuronal cluster. Enrichment of all-neuron DEGs among neuronal cluster markers was analyzed using hypergeometric testing with a background of 2800 genes. Gene ontology (GO) analysis was performed using ToppFun from the ToppGene Suite with default parameters (Chen et al. 2009), and Biological Process GO categories with Benjamini-Hochberg (BH) FDR<0.05 were summarized using REVIGO, with allowed similarity=0.5 and GO term database for *Mus musculus* (Supek et al. 2011). To identify putative Foxp2 direct gene targets, we calculated the Spearman correlation coefficients between *Foxp2* and all other genes in control cells, and then overlapped genes with |ρ|>0.1 and BH FDR<0.05 with the E16.5 brain Foxp2 ChIP long list from (Vernes et al. 2011).

### Gene Expression Omnibus (GEO) accession information

The National Center for Biotechnology Information GEO accession number for the scRNA-seq data reported in this study is GSE130653 (token: mzypiqcotlwtdep).

## Results

### Cortical *Foxp2* deletion impairs reversal learning

We generated cortex-specific *Foxp2* conditional knockout (cKO) mice and control littermates by crossing *Foxp2^flox/flox^* mice (French et al. 2007) with mice expressing *Emx1-* Cre, which induces recombination embryonically in progenitors and projection neurons derived from dorsal telencephalon (Gorski et al. 2002) (Fig. 1A). In adult *Foxp2* cKO mice, we confirmed the absence of Foxp2 protein in the cortex and normal expression in other brain regions (Supplemental Fig. 1A). Consistent with known expression patterns of *Foxp2* in wild-type mouse brain, Foxp2 protein was absent from control hippocampus, where *Emx1-Cre* is also expressed (Ferland et al. 2003; Gorski et al. 2002) (Supplemental Fig. 1A). We observed no gross abnormalities in overall brain morphology of cKO mice (Supplemental Fig. 1A), consistent with a previous study utilizing the same conditional knockout strategy (French et al. 2018). In neonatal frontal cortex, we quantified a >90% reduction in Foxp2 protein content, confirming the efficiency of knockout in developing cortex (Supplemental Fig. 1B).

**Figure 1.**
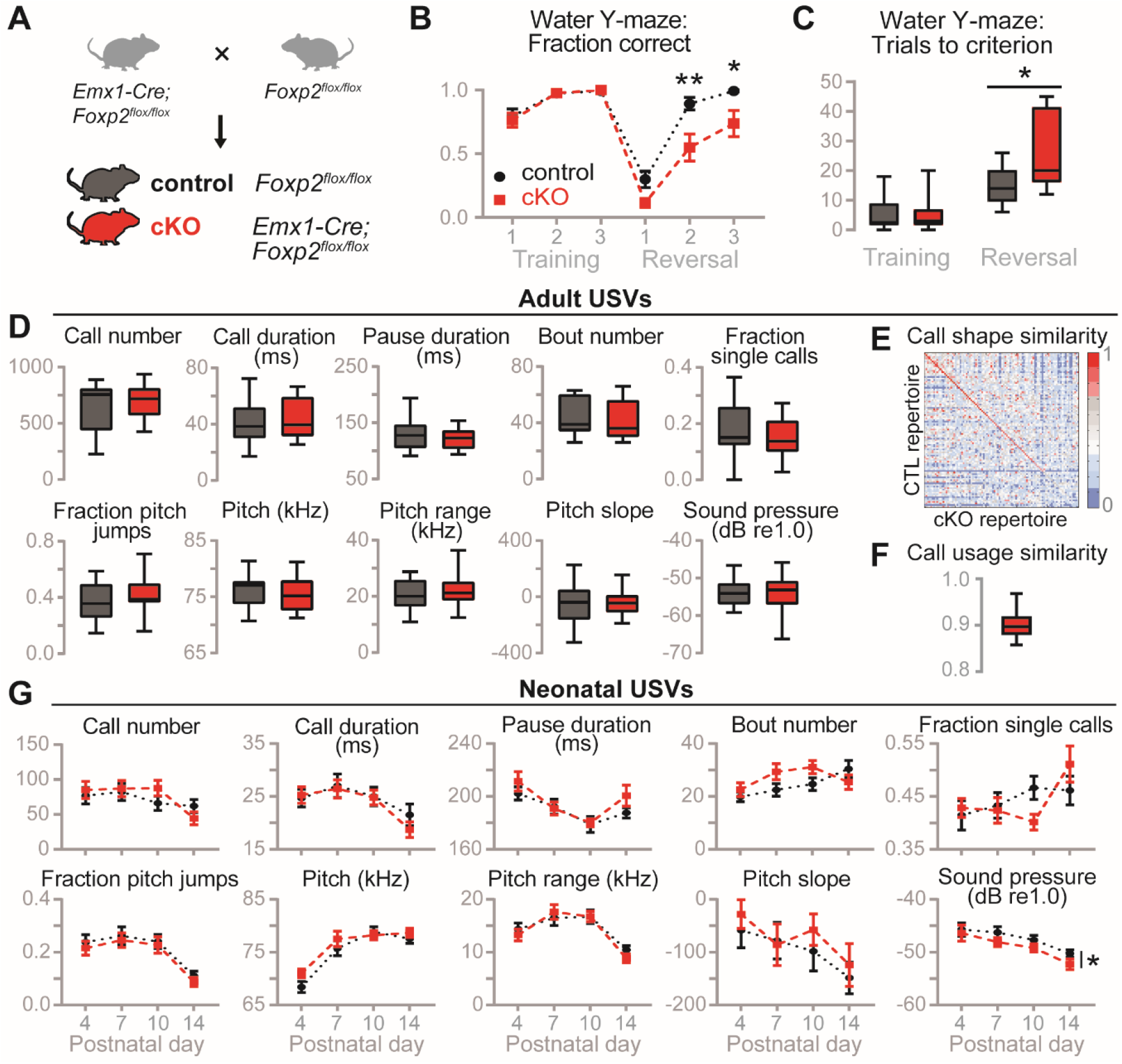
*Foxp2* cKO mice show behavioral inflexibility but normal vocalizations. (A) Breeding scheme to generate control (*Foxp2^flox/flox^*) and *Foxp2* cKO (*Emx1-Cre; Foxp2^flox/flox^*) littermate mice. (B-C) Reversal learning in water Y-maze. *n* = 10-17 per condition. (B) Fraction of correct trials. Data are shown as means (±SEM). (*) *P* < 0.05, (**) *P* < 0.01, two-way ANOVA with Bonferroni’s multiple comparisons test. (C) Number of trials to criterion. Box shows 25-75 percentiles, whiskers show min-max. (*) *P* < 0.05, t-test. (D-G) Analysis of USVs. (D) Adult courtship USVs. Box shows 25-75 percentiles, whiskers show min-max. *n* = 14-15 per condition. (E) Call shape similarity matrix between adult control and cKO USV repertoires (size 100). Scale represents Pearson correlation coefficient. (F) Call usage similarity of adult cKOs compared to controls. Box shows 25-75 percentiles, whiskers show 5-95 percentiles. (G) Neonatal isolation USVs. Data are represented as means (±SEM). (*) *P* < 0.05, two-way ANOVA with Bonferroni’s multiple comparisons test. *n* = 34-42 per condition. Full statistical analysis can be found in Supplemental Table 2.

We evaluated the contribution of cortical *Foxp2* to neurodevelopmental disorder (NDD)-relevant behaviors, such as behavioral flexibility, hyperactivity, anxiety, and social communication. To assess behavioral flexibility in *Foxp2* cKO mice, we utilized a water Y-maze assay and found significant deficits in reversal learning, but not initial acquisition, of escape platform location (Fig. 1B-C, statistics for behavioral testing are provided Supplemental Table 2). As an additional assay of frontal cortical function, we assessed spontaneous alternation in a dry T-maze (Lalonde 2002). While there were no significant differences between genotypes in alternation rate or latency to arm, control mice alternated significantly above chance levels while cKO mice did not (Supplemental Fig. 2A-B, Supplemental Table 2).

We performed additional assays to determine whether locomotor or anxiety phenotypes contributed to *Foxp2* cKO impairment in these cognitive tasks. There were no differences in baseline activity levels in a novel cage (Supplemental Fig. 2C, Supplemental Table 2) or total distance moved in an open field (Supplemental Fig. 2D, Supplemental Table 2). Furthermore, each genotype spent equal amounts of time in the center or open arms of the open field or elevated plus maze, respectively, indicating normal anxiety levels in cKO mice (Supplemental Fig. 2E-F, Supplemental Table 2). Altogether these data indicate that cortical *Foxp2* is required for behavioral flexibility in mice, but not for regulation of activity or anxiety levels.

### Cortical *Foxp2* deletion decreases sound pressure of neonatal vocalizations

We evaluated the contribution of cortical *Foxp2* to social communication by measuring courtship ultrasonic vocalization (USV) production and spectral features in adult *Emx1*-Cre *Foxp2* cKO mice. Using automated call detection methods (Holy and Guo 2005), we found no differences between genotypes in measures related to call number, timing, structure, pitch, or intensity (Fig. 1D, Supplemental Table 2). We next used an automated method to cluster calls into 100 call types (repertoire units or RUs) and compare repertoires between genotypes (Van Segbroeck et al. 2017) (Fig. 1E-F and Supplemental Fig. 2G). This yielded a similarity matrix comparable to matrices generated between cohorts of wild-type C57BL/6 mice (Van Segbroeck et al. 2017), with the top 74 of 100 RUs having Pearson correlations greater than 0.8 (Fig. 1E). Because the similarity matrix does not account for frequency of call types used, we calculated a median (top 50% most used RUs) similarity score of 0.90 and an overall (top 95%) similarity score of 0.86 between control and cKO repertoires (Fig. 1F). Comparing this to the average similarity of 0.91 ± 0.03 between replicate C57BL/6 studies (Van Segbroeck et al. 2017) leads us to conclude that cKO mice do not differ greatly from controls in courtship call structure and usage.

We also investigated the contribution of cortical *Foxp2* to isolation USVs across postnatal development (P4, P7, P10, P14). Again, we found no differences between genotypes in measures related to call number, timing, structure, or pitch (Fig. 1G, Supplemental Table 2). There was, however, a small but significant decrease in the relative sound pressure of calls emitted by *Foxp2* cKO pups across development (Fig. 1G, Supplemental Table 2). This decrease in loudness was not due to somatic weakness, as cKO pups gained weight and performed gross motor functions normally (Supplemental Fig. 2H-J, Supplemental Table 2). In summary, cortical *Foxp2* plays a specific role in loudness of neonatal vocalizations, but not in production or other acoustic features of neonatal or adult vocalizations.

### Cortical *Foxp2* is dispensable for lamination and layer 6 axon targeting

We asked whether abnormalities of corticogenesis could underlie the cognitive deficits in our *Foxp2* cKO mice. Because acute knockdown of *Foxp2* in embryonic cortex was shown to impair neuronal migration (Tsui et al. 2013), we examined cortical layering in cKO mice using DAPI staining of cytoarchitecture and immunohistochemistry for layer markers CUX1 (L2-4), CTIP2 (L5b) and TBR1 (L6). We found no gross abnormalities in layer formation or relative thickness at P7 (Supplemental Fig. 3A-B, statistics for immunohistochemistry are provided Supplemental Table 2). Next, because Foxp2 regulates genes involved in axon outgrowth and guidance in embryonic brain (Vernes et al. 2011), we examined the formation of cortical L6 axon tracts labeled with *golli*-т-eGFP in P14 cKO mice (Jacobs et al. 2007) (Supplemental Fig. 3C). We observed normal formation of L6 axon tracts (including the internal capsule, which contains corticothalamic axons), innervation of thalamic nuclei, and intra-cortical axon and dendrite projections to L4 (Supplemental Fig. 3D). These results confirm other recent findings that *Foxp2* deletion from cortical progenitors and/or neurons does not affect gross cortical layering or targeting of L6 axons (Kast et al. 2019; Medvedeva et al. 2018).

### Cortical *Foxp2* deletion reduces dopamine signaling gene expression

FoxP2 has been shown to regulate expression of the dopamine D1 receptor (D1R) and its downstream effector DARPP-32 in zebra finch striatum (Murugan et al. 2013). Given the long-established link between prefrontal cortical dopamine and behavioral flexibility (Ott and Nieder 2019), we explored the possibility that *Foxp2* deletion impairs reversal learning through dysregulation of cortical dopamine signaling. In cortical L6, Foxp2-expressing corticothalamic projection neurons (CThPNs) are all reported to express DARPP-32, and Foxp2 directly binds the promoter of its gene *Ppp1r1b* in embryonic brain (Hisaoka et al. 2010; Vernes et al. 2011). Retrograde labeling has shown that L6 CThPNs express D1R (Gaspar et al. 1995), and while no studies to date have shown colocalization of Foxp2 and D1R in the cortex, these proteins are highly coexpressed in striatonigral spiny projection neurons (Vernes et al. 2011). Thus, we hypothesized that Foxp2 activates D1R and DARPP-32 expression in mouse L6 CThPNs.

To visualize D1R and DARPP-32 expression, we crossed our *Foxp2* cKO mice with Drd1a-tdTomato BAC reporter mice, which replicate endogenous D1R expression patterns in the cortex, and performed immunohistochemistry at adult and neonatal stages (Ade et al. 2011; Anastasiades et al. 2018). In adult control frontal cortex, DARPP-32 expression closely followed that of Foxp2 in layer 6, but D1R was almost exclusively expressed in Foxp2-negative neurons (Fig. 2B). In agreement with recent studies in Drd1a-tdTomato mouse cortex (Anastasiades et al. 2018), this indicates that mature D1R-expressing neurons are predominantly ITPNs rather than Foxp2/DARPP-32-expressing CThPNs. Unexpectedly, however, cortical *Foxp2* deletion caused a large reduction in D1R-positive cells throughout the adult cortex (Fig. 2B). Thus, cortical Foxp2 is required for proper D1R expression in mature ITPNs.

**Figure 2.**
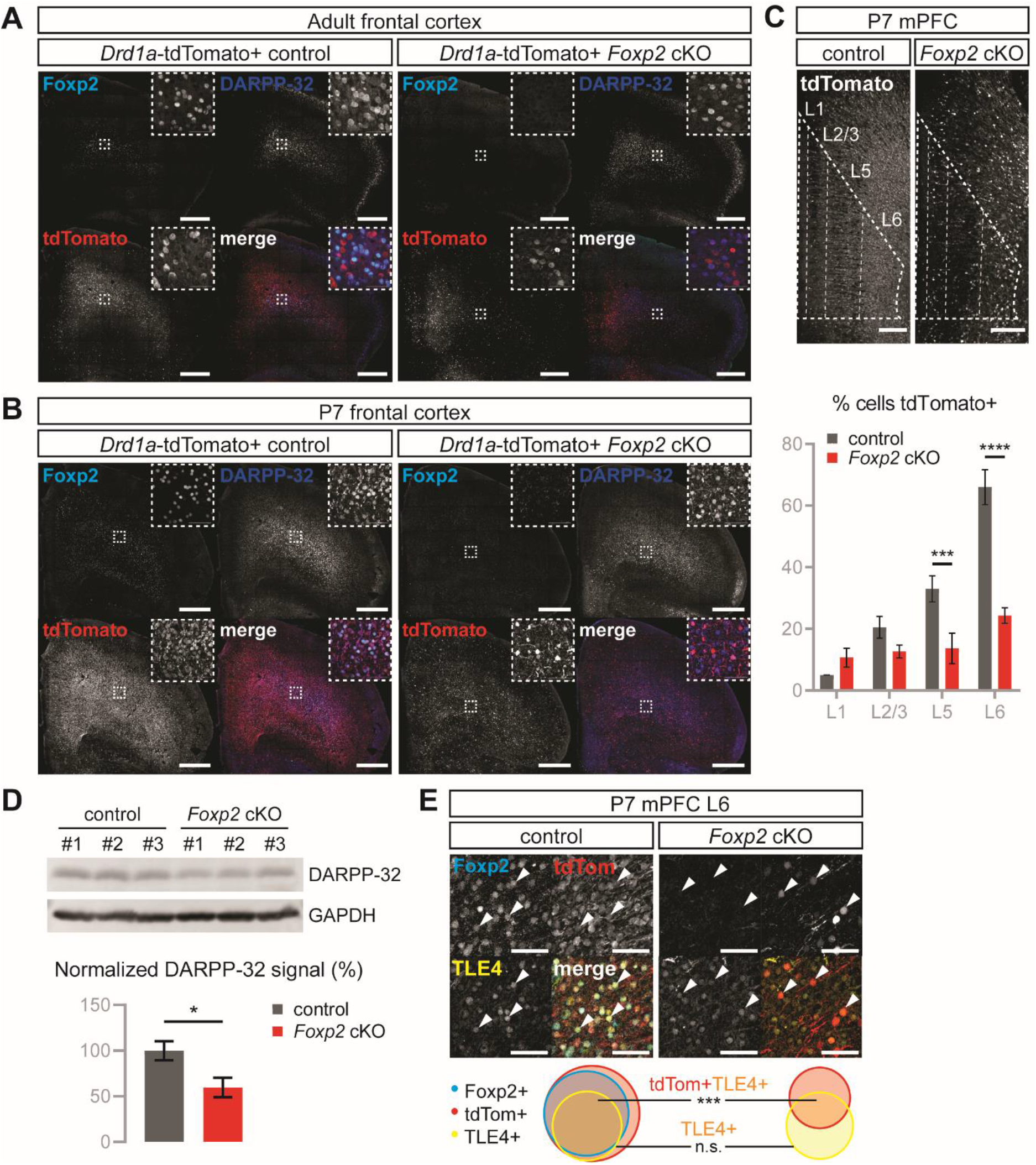
*Foxp2* cKO mice show decreased dopamine signaling proteins in postnatal and adult cortex. (A) IHC for Foxp2, DARPP-32, and signals were normalized*Drdla*-tdTomato in adult control and *Foxp2* cKO frontal cortex. Scale bar: 500 μm. (B) IHC for Foxp2, DARPP-32, and *Drdla*-tdTomato in P7 control and cKO frontal cortex. Scale bar: 500 μm. (C) Top: IHC for *Drdla*-tdTomato in P7 control and cKO mPFC. Scale bar: 200 μm. Bottom: Percentage of DAPI+ cells expressing tdTomato per layer in P7 mPFC. Error bars represent ±SEM. (***) *P* < 0.001, (****) *P* < 0.0001, two-way ANOVA with Bonferroni’s multiple comparisons test. *n* = 2-3 per condition. Full statistical analysis can be found in Supplemental Table 2. (D) Top: Western blot for DARPP-32 and GAPDH loading control from frontal cortical lysates of P7 control and cKO mice. Bottom: Western blot quantification. DARPP-32 signals were normalized to GAPDH signals. Error bars represent ±SEM. (*) *P* < 0.05, t-test. *n* = 3 per condition. (E) Top: IHC for Foxp2, *Drdla*-tdTomato, and TLE4 in P7 control and cKO mPFC L6. Arrowheads indicate cells with protein coexpression. Scale bar: 50 μm. Bottom: Weighted Venn diagrams summarizing Foxp2, *Drdla*-tdTomato, and TLE4 coexpression in P7 control and cKO mPFC L6. (***) *P* < 0.001, t-test. *n* = 2-3 per condition. Full quantification and statistical analysis can be found in Table 1.

In the prefrontal cortex, expression of D1/D1-like receptors is developmentally regulated, with higher expression during earlier stages of development (Andersen et al. 2000; Cullity et al. 2018). Thus, we examined expression of D1R, DARPP-32, and Foxp2 in early postnatal frontal cortex of control and *Foxp2* cKO mice. In contrast with adult control cortex, postnatal control cortex showed a high density of D1R-expressing cells and extensive coexpression of D1R, DARPP-32, and Foxp2 in L6 neurons (Fig. 2B). Again, upon cortical *Foxp2* deletion, we saw a vast reduction of D1R+ cells throughout the frontal cortex (Fig. 2B). Quantification of D1R+ cells by layer in the medial prefrontal cortex (mPFC) revealed significant reductions in L5 (33±3.2% vs. 14±2.8%) and L6 (66±3.9% vs. 25±1.5%), layers where Foxp2 expression normally occurs (Fig. 2C, Supplemental Table 2). In addition, there was a 40% reduction in DARPP-32 protein content in postnatal *Foxp2* cKO cortex (Fig. 2D). These results indicate that in developing cortex, Foxp2 is required for normal expression of dopamine signaling molecules.

To determine whether the decrease in D1R expression in *Foxp2* cKO cortex was due to decreased CThPN density or downregulation of D1R in CThPNs, we examined the CThPN marker TLE4 (Molyneaux et al. 2015) and its coexpression with Foxp2 and D1R in mPFC. In control mice, we found a high degree of overlap among the three proteins in L6 and a moderate degree of overlap in L5 (Fig. 2E, Table 1). In *Foxp2* cKO mice, we saw no change in the percentage of TLE4-positive cells in L5 or L6, but there were significant reductions in TLE4/D1R-positive cells in these layers (Fig. 2E, Table 1). These results agree with recent findings of unaltered neuronal density in L5-6 of mice lacking *Foxp2* through the same conditional knockout strategy (French et al. 2018; Kast et al. 2019). Interestingly, although nearly all (~92%) control TLE4-expressing neurons were Foxp2/D1R-positive, 41% of TLE4+ neurons maintained D1R expression after *Foxp2* deletion (Fig. 2E, Table 1), suggesting that D1R expression is regulated by Foxp2 in only a subset of CThPNs. In summary, we found that postnatal but not adult CThPNs express D1R, and that cortical *Foxp2* is required for proper D1R expression in postnatal CThPNs and adult ITPNs.

### Identification of D1R-expressing cell types in developing frontal cortex

Studies using retrograde labeling and genetic markers have identified excitatory and inhibitory neuronal subtypes expressing D1R in adult mouse mPFC (Anastasiades et al. 2018; Han et al. 2017), but less is known about cell types expressing D1R in developing cortex. To identify these cell types and understand cell type-specific effects of *Foxp2* deletion, we used fluorescence-activated cell sorting (FACS) followed by single-cell RNA-sequencing (scRNA-seq) to genetically profile Drd1a-tdTomato+ frontal cortical neurons from P7 control and *Foxp2* cKO mice (Fig. 3A). Using the 10x Genomics Chromium platform (Zheng et al. 2017), we profiled a total of 7282 cells from 2 mice per genotype, detecting similar numbers of transcripts (median UMI/cell: control=8484, cKO=6832) and genes (median genes/cell: control=2678, cKO=2546) between genotypes (Supplemental Fig. 4A). Using Seurat (Butler et al. 2018) we identified 21 clusters containing cells from all mice examined (Supplemental Fig. 4B-C, Supplemental Table 3), and we annotated cell types by overlapping our cluster marker genes with cluster markers from a published neonatal cortical scRNA-seq dataset (Loo et al. 2019). We identified multiple projection neuron, interneuron, and, unexpectedly, non-neuronal clusters from our Drd1a-tdTomato FACS-scRNA-seq (Supplemental Fig. 4B, D). While *Drd1* was not expressed in every cell, it was expressed in every cluster, and re-clustering *Drd1*+ cells resulted in similar cell types as the full dataset (Supplemental Fig. 4E-F). As *tdTomato* transcripts are roughly double that of *Drd1* in individual cells of Drd1a-tdTomato mouse cortex (Anastasiades et al. 2018), we posited that FACS isolated tdTomato+ cells for which we could not detect *Drd1* transcripts by scRNA-seq. Indeed, using sequence information from the BAC used to generate the Drd1a-tdTomato mice (Ade et al. 2011), we found that *Drd1a*-tdTomato BAC expression is enhanced relative to endogenous *Drd1* expression (Supplemental Fig. 4G). Thus, *Drd1* transcripts appear to be present in both neurons and glia of the developing frontal cortex.

**Figure 3.**
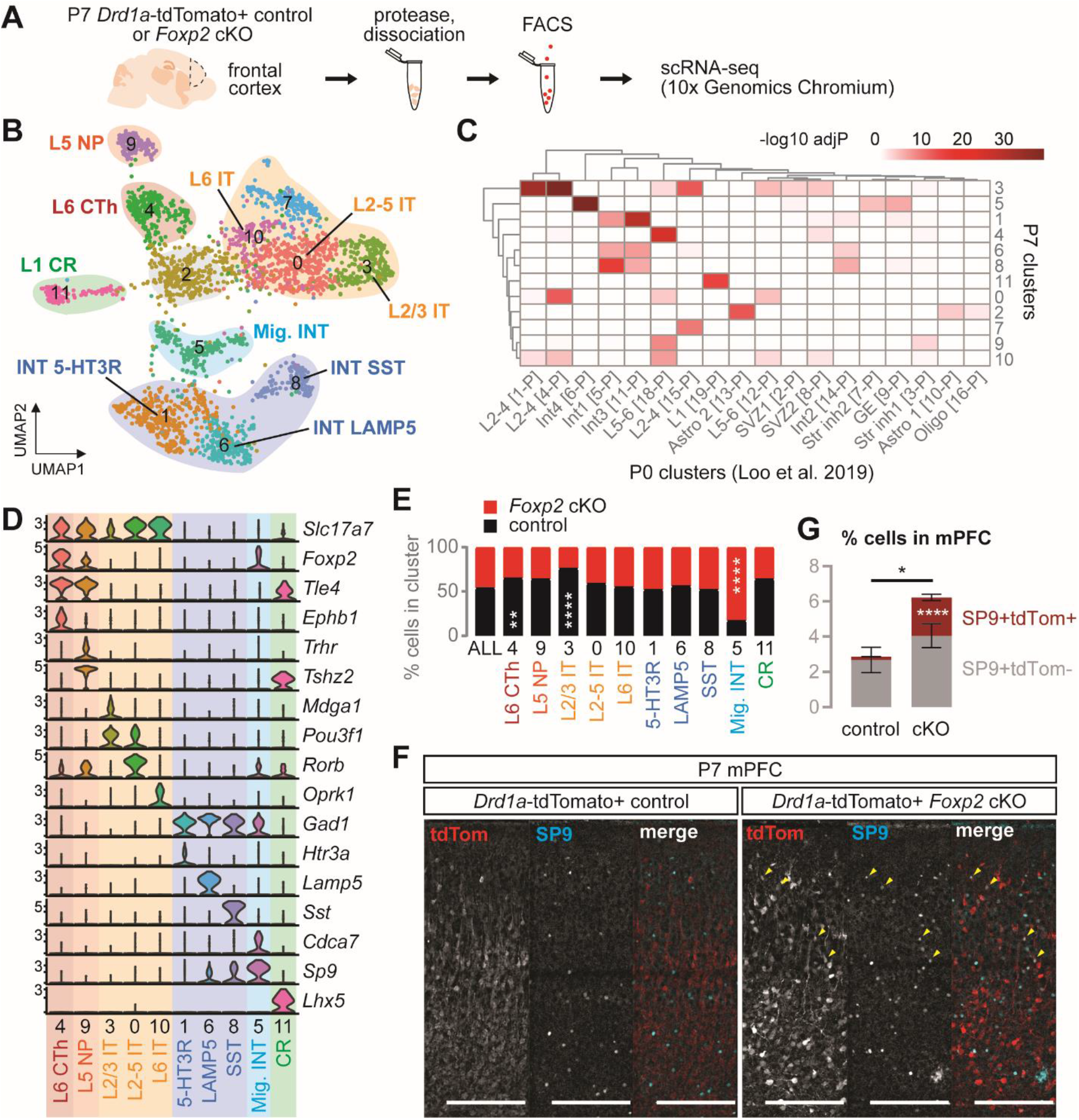
*Foxp2* cKO mice show altered composition of dopamine D1 receptor-expressing neuronal subtypes. (A) Experimental design for D1R scRNA-seq. *n* = 2 per condition. (B) UMAP projection of clusters with neurons combined from both genotypes. (C) Hypergeometric overlaps of neuronal cluster marker genes with P0 mouse cortex cluster marker genes from Loo et al 2019. (D) Violin plots of selected marker genes. Y-axes show log-scaled expression. (E) Percentage of control and cKO cells per cluster. (**) *P* < 0.01, (****) *P* < 0.0001, Fisher’s exact test with Benjamini-Hochberg post-hoc test. (f) IHC for tdTomato and SP9 in P7 control and cKO mPFC. Pial surface is at the top. Arrowheads indicate cells expressing both proteins. Scale bar: 200 μm. (G) Percentage of cells expressing SP9 ± tdTomato in P7 control and cKO mPFC. Data are represented as means±SEM. (*) *P* < 0.05, (****) *P* < 0.0001, t-test. *n* = 3-4 per condition. CR: Cajal-Retzius, CTh: corticothalamic, INT: interneuron, IT: intratelencephalic, Mig INT: migrating interneuron, NP: near-projecting.

**Figure 4.**
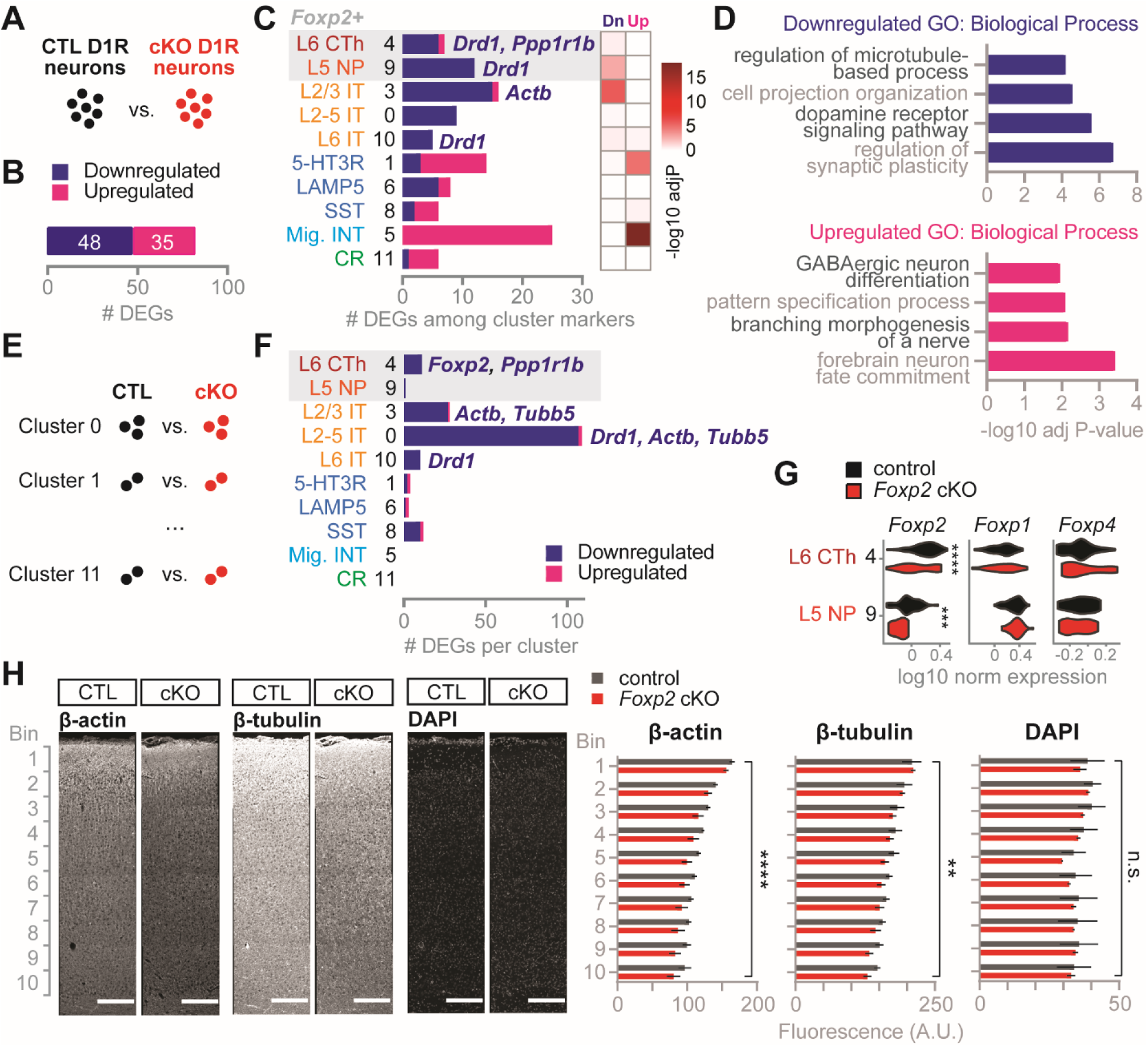
Cortical *Foxp2* deletion induces non-cell-autonomous dysregulation of cytoskeletal genes. (A) Identification of differentially expressed genes (DEGs) between all control and all *Foxp2* cKO neurons from D1R scRNA-seq. (B) Number of DEGs significantly down- or upregulated in cKO neurons. (C) Number (left) and hypergeometric enrichment (right) of DEGs among neuronal cluster marker genes. Gray shaded area indicates clusters with *Foxp2* enrichment. (D) Summarized gene ontology (GO) Biological Process terms for down- and upregulated DEGs. (E) Analysis of DEGs by cluster between control and cKO neurons. (F) Number of DEGs per cluster with selected genes shown. Gray shaded area indicates clusters with *Foxp2* enrichment. (G) Violin plots of *Foxp* gene expression in clusters with *Foxp2* enrichment. (***) *P* < 0.001, (****) *P* < 0.0001, Wilcoxon rank sum test. (H) Left: IHC for β-actin and β-tubulin in P7 control and cKO motor cortex. Right: Quantification of fluorescence intensity averaged by cortical bin. Data are represented as means±SEM. (**) *P* < 0.01, (****) *P* < 0.0001, two-way ANOVA with Bonferroni’s multiple comparisons test. *n* = 3 per condition.

To refine our D1R neuronal subtype classification, we reclustered the neuronal clusters and identified 11 clusters comprised of 2758 cells (Fig. 3B, Supplemental Table 3). Two low-quality clusters (Clusters 2, 7) were excluded from further assessments, and annotation of the remaining clusters based on P0 data revealed multiple subclasses of upper- and lower-layer projection neurons and interneurons (Loo et al. 2019) (Fig. 3C). To delineate projection neuron clusters by their projection specificity, we also examined expression of marker genes from a scRNA-seq dataset with retrograde labeling in adult frontal motor cortex (Tasic et al. 2018) (Fig. 3D). We were able to distinguish L6 CThPNs by *Foxp2, Tle4*, and *Ephb1* (Cluster 4), L5 near-projecting neurons (NPNs) by *Trhr* and *Tshz2* (Cluster 9), L6 ITPNs by *Oprk1* (Cluster 10), and L2/3 ITPNs by *Mdga1* and *Pou3f1* (Cluster 3). Cluster 0 may contain a mix of L2/3 and L5 ITPNs as indicated by expression of both *Pou3f1* and the L5 ITPN marker *Rorb*. Among projection neurons, *Foxp2* expression was restricted to L6 CThPNs and the newly described L5 NPNs, which do not have long-range projections (Tasic et al. 2018) (Fig. 3D). We also distinguished interneuron subtypes in our scRNA-seq data by expression of *Htr3a* (Cluster 1), *Lamp5* (Cluster 6), and Sst (Cluster 8). These results reveal an unprecedented diversity of D1R-expressing neuronal subtypes in the developing frontal cortex, and they identify L5 NPNs as Foxp2-expressing cell types in addition to L6 CThPNs.

### *Foxp2* deletion increases SP9+ interneurons in postnatal cortex

Given that postnatal *Foxp2* cKO mice show reduced D1R expression in CThPNs, but by adulthood show reduced D1R expression in ITPNs, we asked if this potential non-cell-autonomous effect was occurring during development. By examining the proportion of cKO cells in each D1R neuronal cluster, we found significant underrepresentation of cKO cells in L6 CThPN and L2/3 ITPN clusters, and overrepresentation in interneuron Cluster 5 (Fig. 3E). Cluster 5 overlapped significantly with a P0 cluster annotated as migrating cortical interneurons, and these cells expressed high levels of *Cdca7* and *Sp9* while expressing lower levels of mature interneuron subtype markers (Loo et al. 2019) (Fig. 3C-D). Cluster 5 also expressed *Foxp2*, suggesting it arises from an Emx1-negative lineage such as basal forebrain-derived cortical interneurons (Gorski et al. 2002) (Fig. 3D). Immunohistochemistry for SP9 in *Foxp2* cKO mPFC confirmed this increase in total SP9+ cells as well as SP9+D1R+ cells upon *Foxp2* deletion, suggesting the presence of an ectopic population of interneurons in cKO mice (Fig. 3F-G, Supplemental Table 2). Thus, in postnatal frontal cortex, loss of *Foxp2* in Emx1-positive cells causes non-cell-autonomous effects on ITPN D1R expression and cortical interneuron numbers.

### *Foxp2* deletion induces non-cell-autonomous effects on cortical gene expression

To elucidate molecular pathways in cortical D1R neurons affected by *Foxp2* deletion, we performed “pseudo-bulk RNA-seq” differential gene expression analyses between genotypes in our scRNA-seq data. First, we identified differentially expressed genes (DEGs) between all control neurons and all *Foxp2* cKO neurons and found 48 downregulated and 35 upregulated DEGs in cKO neurons (Fig. 4A-B, Supplemental Table 4). In agreement with our immunohistochemistry data, we saw decreased expression of *Foxp2, Drd1*, and *Ppp1r1b* (which encodes DARPP-32) in cKO neurons (Figure 2, Supplemental Table 4). Overlap of these DEGs with our neuronal cluster markers revealed enrichment of downregulated genes in projection neurons and enrichment of upregulated genes in interneurons (Fig. 4C). Summarized gene ontology (GO) terms associated with downregulated DEGs indicated abnormal synaptic plasticity, dopamine signaling, projection organization, and microtubule-based processes in cKO neurons (Fig. 4D, Supplemental Table 4). Summarized GO terms for upregulated DEGs were consistent with and likely driven by the ectopic immature interneuron Cluster 5, which was comprised of more cKO neurons than control neurons (Fig. 3E-G, Fig. 4D, Supplemental Table 4).

In a differential gene expression approach less driven by imbalanced cell type proportions between genotypes, we also identified DEGs within each neuronal cluster (Fig. 4E, Supplemental Table 5). L6 CThPNs and L5 NPNs in cKO mice showed surprisingly few gene expression changes, but we confirmed *Ppp1r1b* as a CThPN-specific downregulated DEG (Fig. 4F). This relatively small number of DEGs in *Foxp2-* expressing neurons was not due to upregulation of related genes *Foxp1* or *Foxp4* (Fig. 4G). In contrast, L2-5 ITPN clusters showed a greater number of DEGs, including downregulation of genes encoding the cytoskeletal proteins β-actin (*Actb*) and β-tubulin (*Tubb5*) (Fig. 4F). To confirm these decreases, we performed immunohistochemistry for β-actin and β-tubulin in P7 control and *Foxp2* cKO cortex and saw decreased expression of both proteins across the cortical mantle (Fig. 4H, Supplemental Table 2). Thus, while *Foxp2* expression is restricted to deep-layer non-ITPNs, its deletion causes non-cell-autonomous changes including cytoskeletal gene downregulation in upper-layer ITPNs.

Next, to overcome the differential sampling of cell types between genotypes due to Foxp2 regulation of Drd1a-tdTomato, we generated an independent FACS-scRNA-seq dataset from mice expressing the *golli*-т-eGFP (GTE) reporter, which is expressed in control Foxp2-positive CThPNs but not decreased in cKO cortex (Supplemental Fig. 5A-B). Using similar analysis methods as with the Drd1a-tdTomato dataset, we identified projection neuron clusters and calculated pseudo-bulk RNA-seq DEGs between genotypes, 24 of which were downregulated in cKO and 23 upregulated (Supplemental Fig. 5C-F, Supplemental Table S6). Summarized GO terms associated with downregulated DEGs were related to neuronal projection organization and synaptic signaling, similar to the downregulated GO terms in the Drd1a-tdTomato dataset (Supplemental Fig. 5G, Supplemental Table S6). Notably, *Actb* appeared among the downregulated DEGs in both datasets (Supplemental Fig. 5H, Supplemental Table S6). In summary, cortical *Foxp2* deletion induces both cell-autonomous and non-cell-autonomous decreases in dopamine-related, synaptic, and projection-related gene expression.

### Putative direct targets of Foxp2 in postnatal cortex

In another approach to identify Foxp2 targets in a manner uninfluenced by altered Drd1a-tdTomato expression in *Foxp2* cKO cortex, we searched for genes correlated (i.e. activated targets) or anticorrelated (i.e. repressed targets) with *Foxp2* expression across control neurons in the *Drd1a*-tdTomato dataset (Fig. 5A, Supplemental Table 7). To identify potential direct targets, we overlapped these genes with targets from a Foxp2 promoter-binding assay performed in embryonic mouse brain (Vernes et al. 2011). Then, to determine their cell type-specificity, we overlapped these putative direct Foxp2 targets with the Drd1a-tdTomato neuronal cluster markers (Fig. 5B). We found shared and distinct Foxp2-activated targets between CThPNs and NPNs, several of which may exert non-cell-autonomous effects through extracellular matrix organization (*Col23a1, P4ha1*), cell-cell signaling (*Islr2, Plxna2, Sdk1*), or synaptic activity (*Cacna1a, Calm2, Cbln1, Grm3, Lrrtm2*) (Fig. 5B). We also identified Foxp2-repressed targets, which were markers for ITPNs, interneurons, and CR cells, suggesting that Foxp2 plays a role in repressing these identities in CThPNs and NPNs (Fig. 5B). These results provide potential mechanisms by which Foxp2 may exert non-cell-autonomous effects and contribute to the maintenance of deep-layer cortical projection neuron identity.

**Figure 5.**
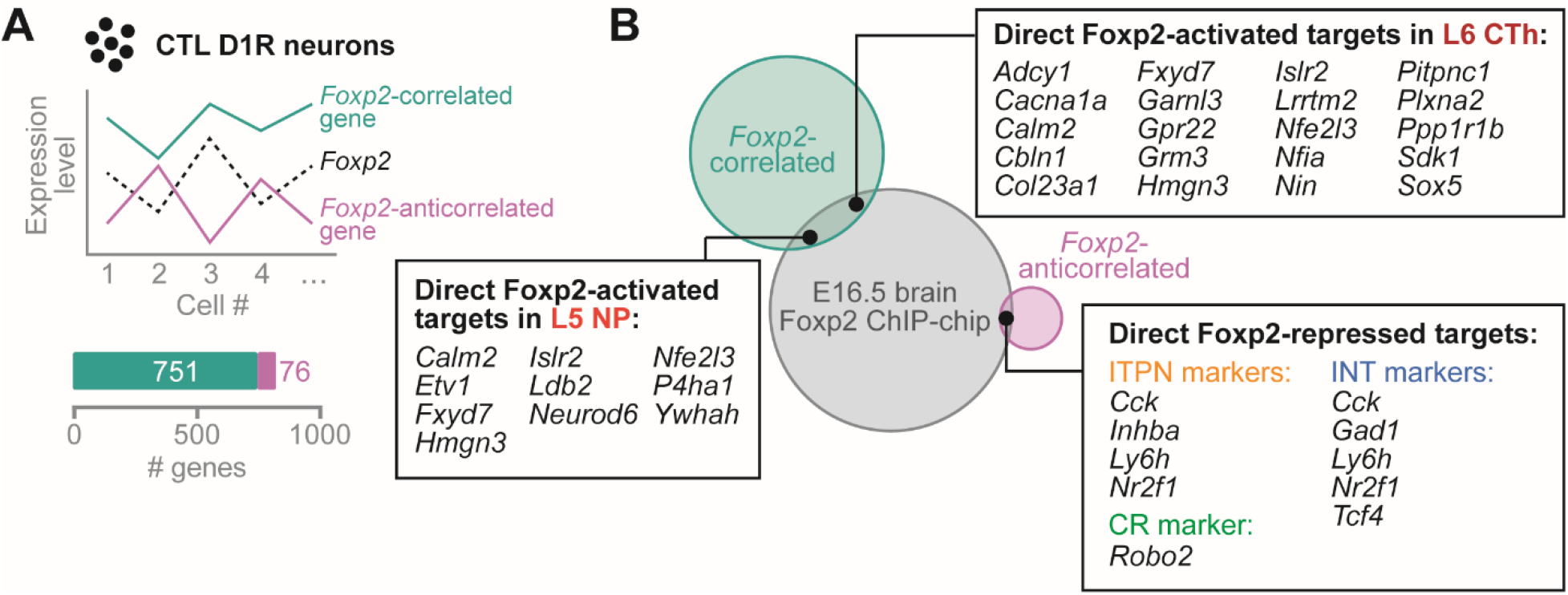
Identification of genes directly regulated by Foxp2 in the cortex. (A) Top: Analysis of genes correlated or anticorrelated with *Foxp2* expression across control neurons from D1R scRNA-seq. Bottom: Number of Foxp2-correlated or -anticorrelated genes. (B) Overlap of Foxp2-correlated or -anticorrelated genes with embryonic brain Foxp2 ChIP-chip targets from (Vernes et al. 2011) and with neuronal cluster marker genes. CR: Cajal-Retzius, CTh: corticothalamic, INT: interneuron, IT: intratelencephalic, Mig INT: migrating interneuron, NP: near-projecting.

## Discussion

### *Foxp2*-regulated cortical dopamine signaling and behavioral flexibility

We have demonstrated specific roles for cortical *Foxp2* in reversal learning, a form of behavioral flexibility, and cortical dopamine D1R signaling throughout the postnatal lifespan (Fig. 6). Cortical dopamine signaling regulates many cognitive functions, including behavioral flexibility (Floresco 2013; Ott and Nieder 2019), and specific manipulations of cortical D1R signaling or D1R-D2R interactions can modulate reversal learning ability (Calaminus and Hauber 2008; Mizoguchi et al. 2010; Thompson et al. 2016). Thus, we suggest that the reversal learning deficits in *Foxp2* cKO mice arise from their decreased expression of cortical dopamine D1 receptors. Other studies linking *Foxp2* to cognitive function and dopamine signaling found that humanized *Foxp2* mice demonstrate enhanced strategy set-shifting, another form of behavioral flexibility, and altered frontal cortical dopamine concentrations (Enard et al. 2009; Schreiweis et al. 2014). Furthermore, specific knockdown of *Drd1* in the prefrontal cortex impairs strategy set-shifting (Cui et al. 2018). Interestingly, we only observed significant deficits in cognitive function in the water Y-maze but not in the dry T-maze (Fig 1B-C, Supplemental Fig. 2A). Given that dopamine release in the frontal cortex is influenced by acute stress (Arnsten 2009), the potential D1R-mediated cognitive impairments in *Foxp2* cKO mice could be exacerbated in aversive tasks such as the water Y-maze.

**Figure 6.**
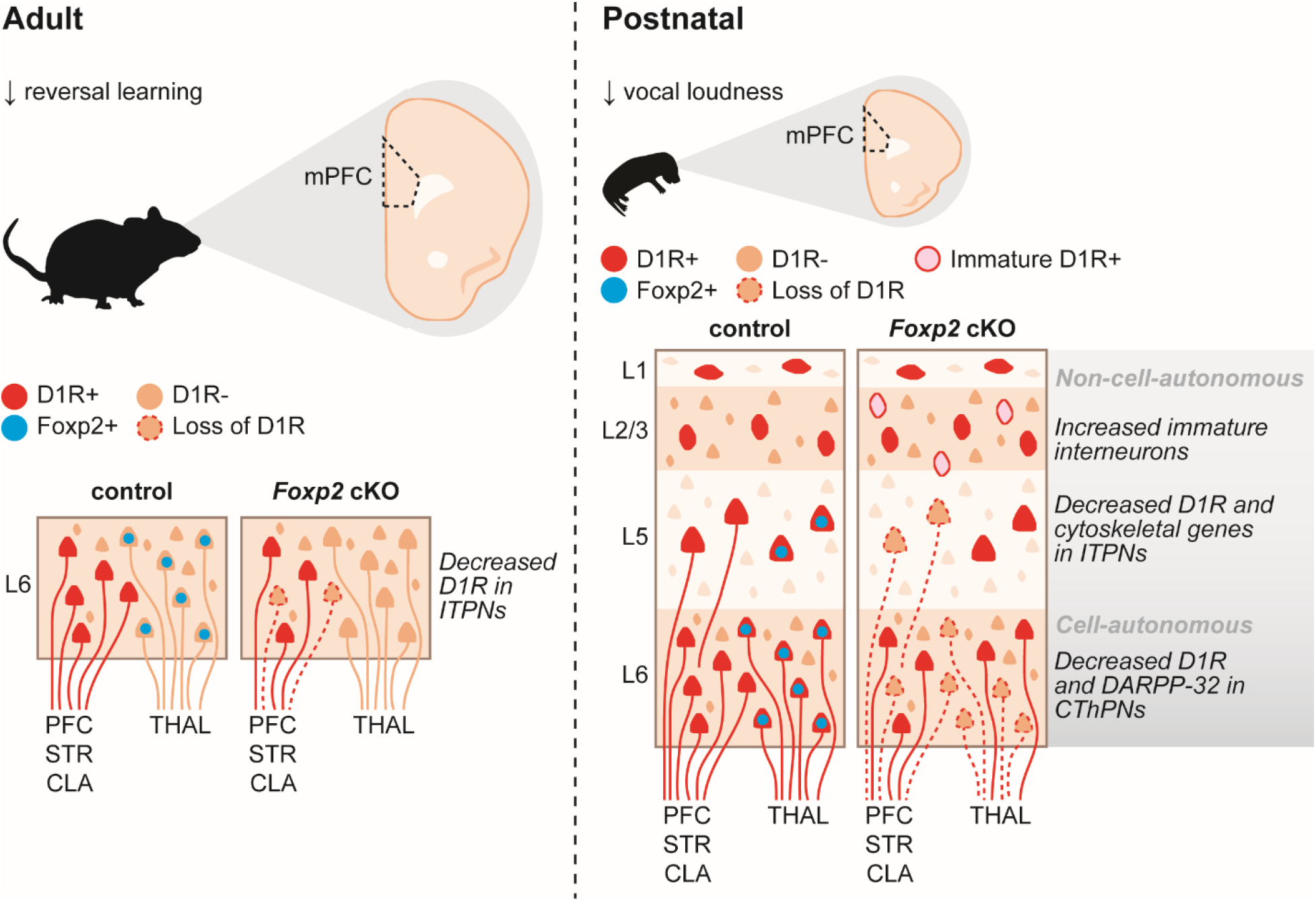
Summary of behavioral and molecular findings in *Foxp2* cKO mice. CLA: claustrum, CThPNs: corticothalamic projection neurons, D1R: dopamine D1 receptor, ITPNs: intratelencephalic projection neurons, L: layer, mPFC: medial prefrontal cortex, STR: striatum, THAL: thalamus.

Cortical *Foxp2* may mediate flexible behaviors through multiple circuit pathways in the brain. Recent optogenetic experiments have demonstrated involvement of both corticothalamic and corticostriatal neurons in probabilistic reversal learning (Nakayama et al. 2018). Thus, the reversal learning deficits in *Foxp2* cKO mice may be due to dysregulation of *Drd1* or other genes in CThPNs and ITPNs (Fig. 4), the latter of which encompass corticostriatal neurons. In addition, the hippocampus is known to have a prominent role in reversal learning in rodents and humans (Mala et al. 2015; Vila-Ballo et al. 2017). While Foxp2 protein has limited expression in the hippocampus (Supplemental Fig. 1), we cannot rule out the possibility that loss of *Foxp2* in the cortex might ultimately affect hippocampal function through disruption to brain circuitry.

### Genetic diversity of dopaminoceptive cell types in developing cortex

Studies spanning over two decades, reviewed in (Anastasiades et al. 2018), have identified diverse neuronal subtypes expressing dopamine D1 receptors in the adult cortex. However, much less is known about cell types expressing D1R in the developing cortex, despite reports of postnatal regulation of D1R expression in this region (Andersen et al. 2000; Brenhouse et al. 2008; Cullity et al. 2018; Tarazi et al. 1999). To better understand the effects of *Foxp2* deletion on developing D1R neuronal subtypes, we conducted the first cell type characterizations of *Drd1a*-tdTomato postnatal frontal cortex (Ade et al. 2011). A recent study of these mice in adulthood found extensive expression of D1R in ITPNs but limited expression in CThPNs (Anastasiades et al. 2018), a finding corroborated by the absence of D1R in adult Foxp2-expressing CThPNs (Fig. 2A). In stark contrast, we found a high degree of D1R expression in both CThPNs and ITPNs of postnatal frontal cortex via immunostaining and single-cell transcriptomics (Fig. 2E, Fig. 3B). Similarly, both adult and postnatal cortex show D1R expression in 5HT3R and calretinin-expressing (L1 CR) interneuron subtypes, but only postnatal cortex shows D1R expression in somatostatin (SST)-positive interneurons (Anastasiades et al. 2018) (Fig. 3B). These results suggest transient expression of D1R in certain cell types of postnatal frontal cortex, such as CThPNs and SST interneurons. Additionally, we identified glial expression of *Drd1* in our scRNA-seq data, supporting evidence for the presence of D1-like receptors in prefrontal cortical astrocytes (Liu et al. 2009; Vincent et al. 1993) (Supplemental Fig. 4). The developmental functions of these early D1R expression patterns in neurons and potentially glia remain an interesting area of study to be elucidated.

### *Foxp2* regulation of interneuron development in the cortex

In addition to changes in dopamine gene expression, we found that cortical *Foxp2* deletion produced an ectopic population of D1R-expressing interneurons (Fig. 3E-G). These cells closely matched the gene expression signature of ganglionic eminence-derived migrating interneurons identified by cell type profiling of neonatal mouse cortex (Loo et al. 2019) (Fig. 3C). Control of interneuron migration by cortical *Foxp2* could explain the abnormal cell migration seen after *Foxp2* knockdown at E13/14, when *Foxp2*-expressing projection neurons have already migrated to the cortical plate and are positioned to signal to incoming interneurons (Tsui et al. 2013). Intriguingly, Emx1-Cre-mediated cortical deletion of *Satb2*, another projection neuron-specific transcription factor, was also recently shown to influence the migration and connectivity of ganglionic eminence-derived interneurons in the cortex (Wester et al. 2019). Our results provide further evidence that cortical interneuron development depends on proper development and function of projection neurons.

Multiple mechanisms could contribute to excess migrating interneurons in *Foxp2* cKO cortex. Altered dopamine D1 signaling itself could promote over-migration of interneurons into the cortex from the ganglionic eminences, as pharmacological D1R modulation has been shown to guide interneuron migration in embryonic forebrain slice preparations (Crandall et al. 2007). Alternatively, altered dopamine sensitivity or cytoskeletal function of projection neurons may prevent activity-dependent apoptosis of early interneurons, which normally occurs around the postnatal time point examined in our study (Lim et al. 2018). Integration of these excess interneurons into frontal cortical circuits could then impair circuit function and lead to behavioral deficits. Indeed, 22q11 deletion and *Plaur* mouse models of ASD also demonstrate abnormal interneuron number and positioning as well as reversal learning deficits (Bissonette et al. 2010; Meechan et al. 2013).

### Limited roles of cortical *Foxp2* in mouse vocalization

Mice with *Foxp2* mutations commonly exhibit USV abnormalities (Castellucci et al. 2016; Chabout et al. 2016; French and Fisher 2014; Gaub et al. 2016), which have been attributed to its functions in the striatum (Chen et al. 2016), cerebellum (Fujita-Jimbo and Momoi 2014; Usui et al. 2017b), and laryngeal cartilage (Xu et al. 2018). Cortical *Foxp2* deletion using *Nex-Cre* was previously shown to alter adult USVs in a social context-dependent manner (Medvedeva et al. 2018), but in our study, Emx1-Cre-mediated deletion did not appear to impact adult courtship USVs (Fig. 1D-F). Several methodological differences may account for these discrepancies. Nex-Cre causes recombination around embryonic day (E) 11.5 in postmitotic projection neurons, whereas Emx1-Cre acts by E10.5 in both projection neurons and their progenitors (Goebbels et al. 2006; Gorski et al. 2002). Thus, perhaps earlier deletion of *Foxp2* from the cortex induces developmental compensation in vocalization circuitry that cannot occur after postmitotic neuronal deletion. Another possibility is that the superovulated females used to elicit courtship calls in the previous study exposed differences between genotypes that we could not detect using females in a natural ovulation state. Furthermore, our analysis did not parse calls based on duration, direction, or size of pitch jumps as did the previous study, but our call repertoire analysis suggested high overall similarity between control and cKO vocalizations.

Neonatal isolation USVs were also largely unaffected by loss of cortical *Foxp2*, contrasting with the USV reductions in pups with cortical *Foxp1* deletion or cerebellar *Foxp2* knockdown (Usui et al. 2017a; Usui et al. 2017b) (Fig. 1G). Whereas Foxp2 is expressed in CThPNs, Foxp1 is likely expressed in callosal and corticostriatal neurons based on its coexpression with Satb2 (Hisaoka et al. 2010; Sohur et al. 2014; Sorensen et al. 2015). Furthermore, cortical layering is altered in *Foxp1* cKO but not *Foxp2* cKO mice (Usui et al. 2017a). Thus, proper positioning and function of ITPNs and cerebellar output neurons may be more essential to USV production than CThPNs.

Cortical *Foxp2* deletion did decrease the sound pressure of USVs across postnatal development (Fig. 1G). Homozygous *Foxp2-R552H* mutant pups also emit quieter ultrasonic distress calls, which correlates with overall developmental delay of mutants (Gaub et al. 2010; Groszer et al. 2008). However, cortical *Foxp2* alone at least partly contributes to modulation of call loudness, as our cKO pups showed grossly normal development (Supplemental Fig. 2H-J). By adulthood, heterozygous *Foxp2-R552H* mutants emit abnormally loud courtship USVs and show ectopic positioning of L5 laryngeal motor cortex neurons (Chabout et al. 2016; Gaub et al. 2016). Whether this neuronal population is altered in *Foxp2* cKO pups and contributes to call loudness remains to be explored. As to the possible contribution of dopamine D1 signaling to call loudness, very few studies have explored this area. Systemic D1-like receptor blockade during rat development has been shown to increase USV sound pressure at later postnatal ages, while blockade in adult rats recapitulates the abnormal laryngeal neurophysiology seen in Parkinson’s-related hypophonia (Cuomo et al. 1987; Feng et al. 2009). Whether USV loudness is modulated specifically by cortical D1Rs remains to be determined and would clarify the mechanisms by which *Foxp2* cKO pups emit quieter calls.

### Neurodevelopmental disorder gene regulation by Foxp2 in the cortex

Foxp2 regulation of cortical dopamine signaling may inform our understanding and treatment of NDDs. *FOXP2* variation has recently been associated with ASD and ADHD (Demontis et al. 2019; Reuter et al. 2017; Satterstrom et al. 2019), and genetic perturbations of dopamine signaling have also been implicated in NDDs (Money and Stanwood 2013). *Foxp2* cKO mice phenocopy the decreased cortical dopamine gene expression (*Drd1, Ppp1r1b*) and reversal learning impairments seen in 16p11.2 and *Tbr1* mouse models of ASD (Huang et al. 2014; Portmann et al. 2014; Yang et al. 2015), suggesting a possible convergent phenotype of dysregulated cortical dopamine signaling in NDDs affecting behavioral flexibility.

Several other Foxp2-regulated genes are of interest due to their connection with NDDs. Overlap of direct Foxp2 targets with DEGs from *Tbr1* models of ASD reveals potential co-activated (*Adcy1, Cbln1, Grm3, Lrrtm2, Nfe2l3, Nfia, Nin, Sdk1, Ppp1r1b, Sox5*) and co-repressed (*Inhba*) genes by Foxp2-TBR1 interaction in the cortex (Bedogni et al. 2010; Deriziotis et al. 2014; Fazel Darbandi et al. 2018; Vernes et al. 2011). In addition, *Foxp2* expression was anticorrelated with the ASD gene *Mef2c* (Supplemental Table 7), which is directly repressed by Foxp2 in the striatum to control wiring of cortical synaptic inputs (Chen et al. 2016). Furthermore, recently identified ADHD-associated loci include *Foxp2* as well as the Foxp2-correlated genes *Dusp6, Pcdh7*, and *Sema6d*, the last of which is a direct Foxp2 target in the brain (Demontis et al. 2019; Vernes et al. 2011) (Supplemental Table 7). We note that our direct target analysis is limited to Foxp2-bound promoters in embryonic brain (Vernes et al. 2011), and recent evidence indicates that FOXP2 promotes chromatin accessibility at enhancers to regulate gene expression (Hickey et al. 2019). Thus, genome-wide targets of Foxp2 must be identified at various developmental stages for a full understanding of its functions in the cortex, including the molecular mechanism of its regulation of D1R expression.

In summary, our work on cortical *Foxp2* represents a step forward in elucidating neural mechanisms underlying cognition and vocal communication. Importantly, we identified dysregulated molecular pathways upon cortical *Foxp2* deletion in the absence of general cortical development abnormalities. Moreover, these findings provide insights into the etiology and treatment of FOXP2-related and other neurodevelopmental disorders affecting behavioral flexibility.

## Acknowledgments

Our sincerest thanks to: Peter Tsai, Maria Chahrour, Todd Roberts, Jane Johnson, Ashley Anderson, and Ana Ortiz for providing feedback on the manuscript; Dr. Shari Birnbaum at the UT Southwestern Rodent Behavior Core for collecting behavior data and contributing helpful advice; and Dr. Bernd Gloss for providing BAC sequence information for the Drd1a-tdTomato scRNA-seq analysis. We also thank the Neuroscience Microscopy, Whole Brain Microscopy, and Flow Cytometry Facilities at UT Southwestern. G.K. is a Jon Heighten Scholar in Autism Research at UT Southwestern.

## Author Contributions

M.C. and G.K. designed the study. M.C. collected and analyzed behavior, immunohistochemistry, and scRNA-seq data, and wrote the manuscript. S.L.H. and A.K. performed bioinformatic analyses on scRNA-seq data. M.H. collected behavior data, maintained mouse lines, and performed genotyping.

