## Supplemental Material for "Cortical *Foxp2* supports behavioral flexibility and developmental dopamine D1 receptor expression"

### Supplemental Tables

Supplemental Table 1. PCR genotyping primers for mouse lines used in this study.

Supplemental Table 2. Full statistics for behavior and immunohistochemistry data.

Supplemental Table 3. scRNA-seq cluster marker genes for FACS-isolated *Drd1a*-tdTomato+ cells and reclustered neurons.

Supplemental Table 4. Pseudo-bulk RNA-seq DEGs between genotypes in all *Drd1a*-tdTomato+ neurons and summarized Biological Process GO categories.

Supplemental Table 5. Pseudo-bulk RNA-seq DEGs between genotypes by neuronal cluster in *Drd1a*-tdTomato+ neurons.

Supplemental Table 6. scRNA-seq analyses for FACS-isolated *golli*-τ-eGFP+ neurons.

Supplemental Table 7. Spearman correlations between *Foxp2* and all other genes in control *Drd1a*-tdTomato+ neurons.

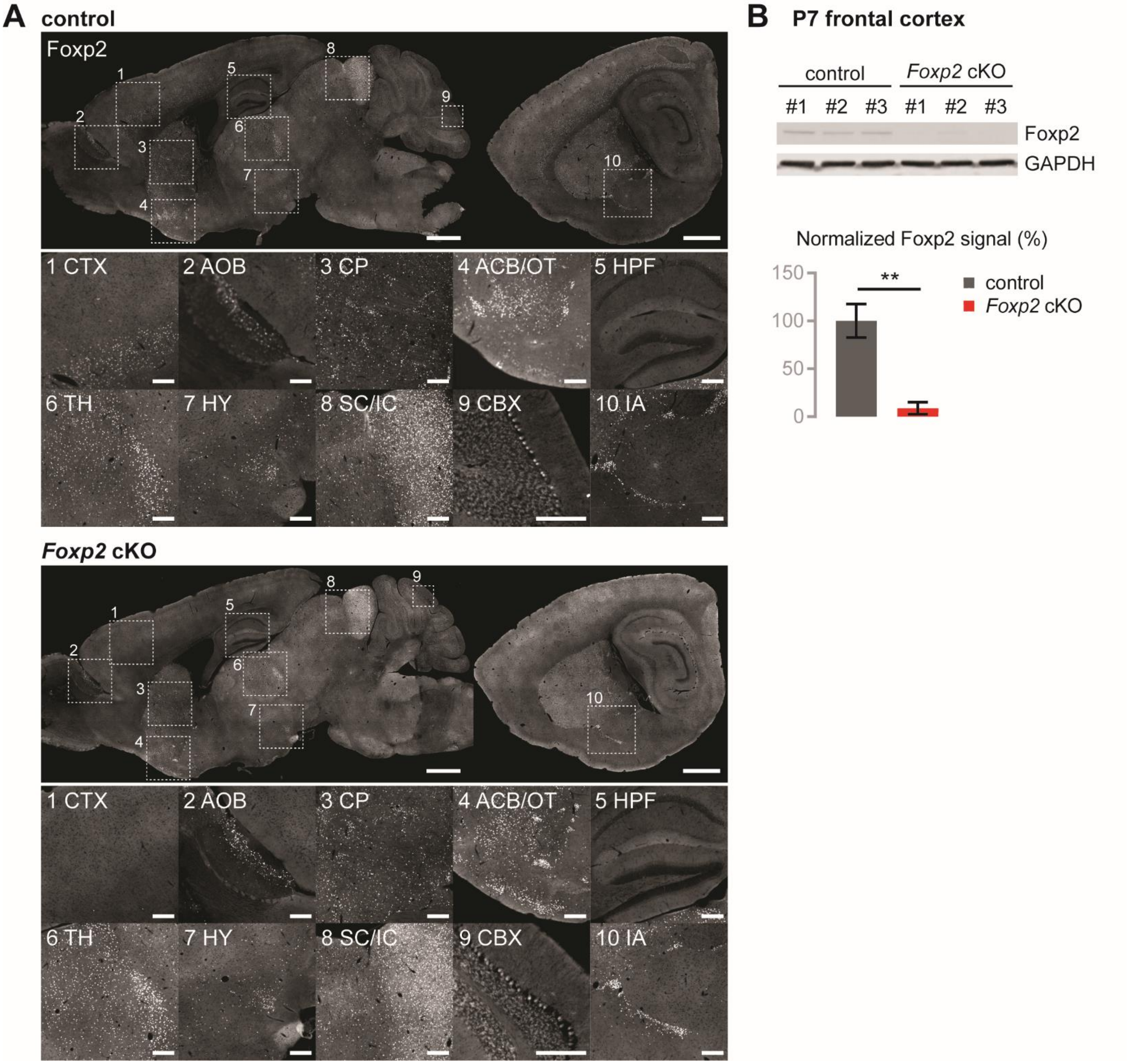


**Supplemental Figure 1. Cortex-specific deletion of *Foxp2* by *Emx1*-Cre.** (A) IHC for Foxp2 in sagittal brain sections from adult control (*Foxp2*$^{flox/flox}$) and *Foxp2* cKO (*Emx1-Cre; Foxp2*$^{flox/flox}$) mice. Top scale bar: 1000 µm, bottom scale bars: 200 µm. (B) Top: Western blot for Foxp2 and GAPDH from frontal cortical lysates of P7 control and cKO mice. Bottom: Western blot quantification. Foxp2 signals were normalized to GAPDH signals. Error bars represent ±SEM. (**) $P < 0.01$, t-test. $n = 3$ per condition. ACB: nucleus accumbens, AOB: accessory olfactory bulb, CBX: cerebellar cortex, CP: caudoputamen, CTX: cortex, HPF: hippocampal formation, HY: hypothalamus, IA: interposed amygdalar nucleus, IC: inferior colliculus, OT: olfactory tubercle, SC: superior colliculus, TH: thalamus.

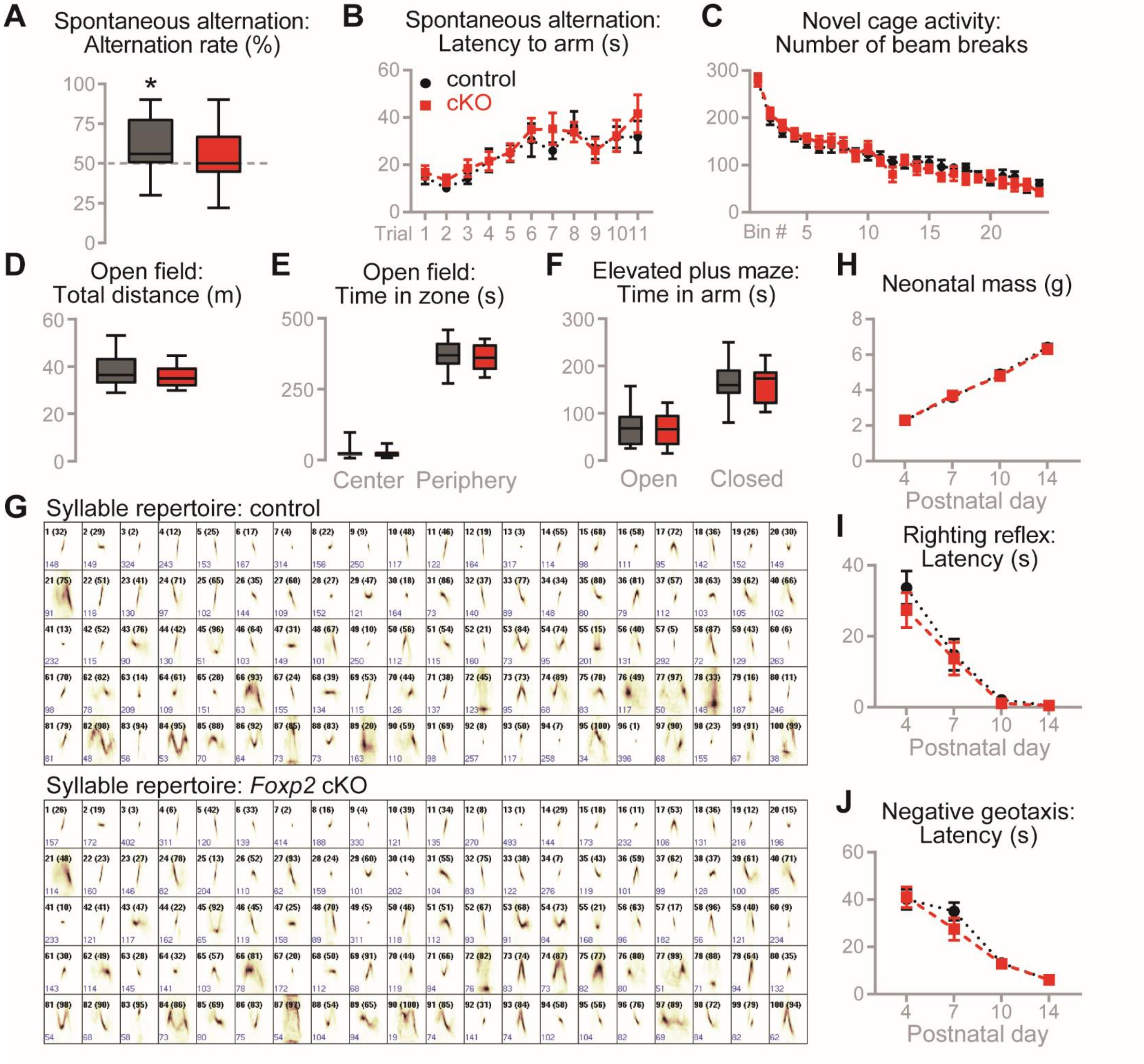


**Supplemental Figure 2. Additional behavioral assessments in *Foxp2* cKO mice.** (A-B) Spontaneous alternation in T-maze. *n* = 16-19 per condition. (A) Spontaneous alternation rate. Box shows 25-75 percentiles, whiskers show min-max. (*) $P < 0.05$ compared to chance levels (50%), t-test. (B) Latency to arm. Data are represented as means (±SEM). (C-F) Activity and anxiety measures. *n* = 13 per condition. (C) Number of infrared beam breaks in a novel cage per 5-minute bin. Data are represented as means (±SEM). (D) Total distance moved in open field. Box shows 25-75 percentiles, whiskers show min-max. (E) Time in zone in open field. Box shows 25-75 percentiles, whiskers show min-max. (F) Time in zone in elevated plus maze. Box shows 25-75 percentiles, whiskers show min-max. (G) Call shapes for control and *Foxp2* cKO repertoires (size 100). (H-J) Neonatal development assessments. Data are represented as means (±SEM). *n* = 18-23 per condition. (H) Neonatal mass in grams. (I) Righting reflex latency. (J) Negative geotaxis latency. Full statistical analysis can be found in Supplemental Table 2.

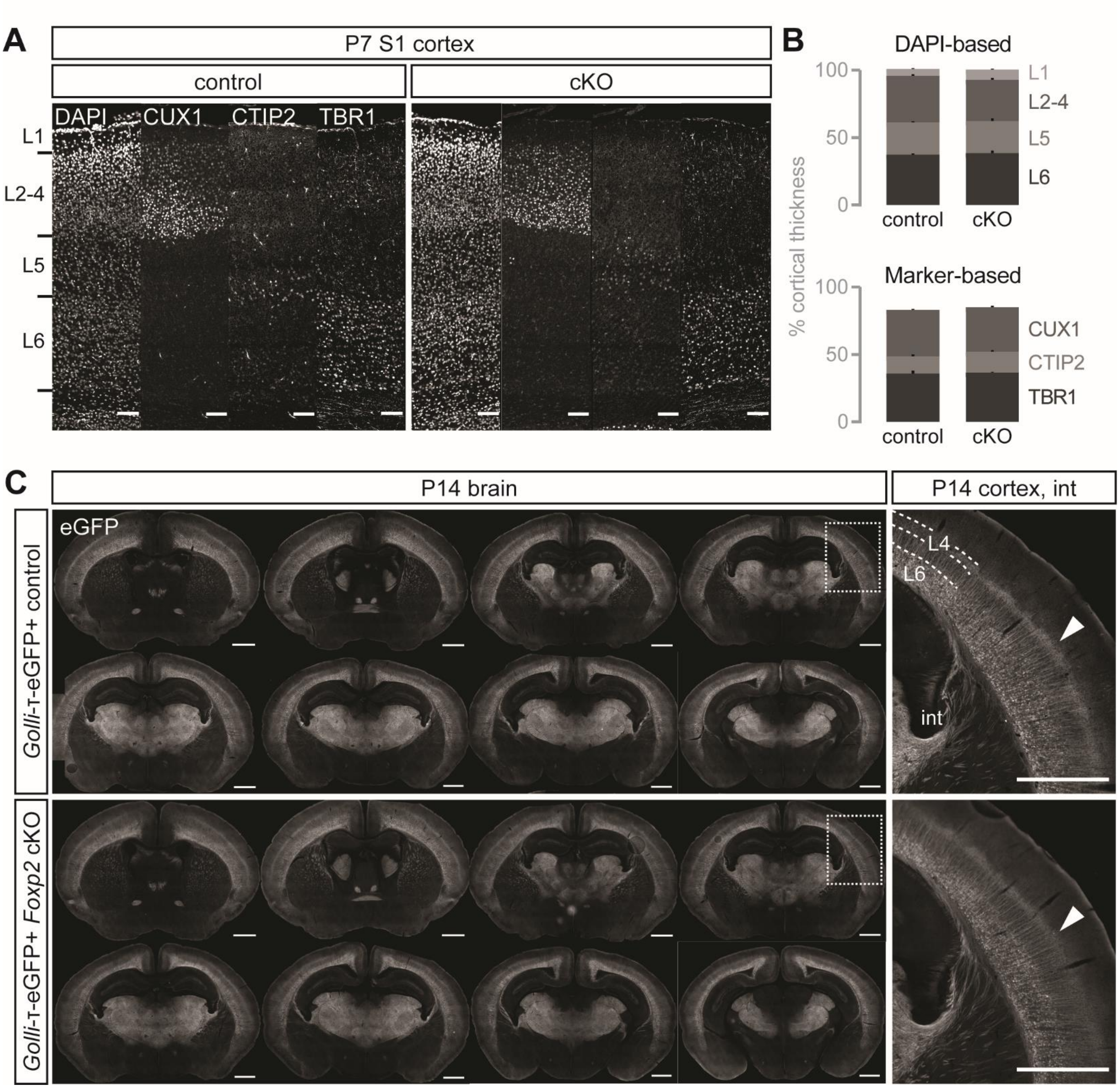


**Supplemental Figure 3. *Foxp2* cKO mice show normal gross cortical development.** (A) IHC for cortical layer markers in P7 control and cKO primary somatosensory (S1) cortex. Scale bar: 100 µm. (B) Quantification of relative layer thickness based on DAPI cytoarchitecture (top) and layer markers (bottom). Data are represented as means (±SEM). $n$ = 3 per condition. Full statistical analysis can be found in Supplemental Table 2. (C) IHC for eGFP in P14 *golli*-τ-eGFP+ control and cKO brain sections (left) and corticothalamic axons (right). Arrowheads indicate L6 axon and dendrite terminations in L4. Int: internal capsule. Scale bars: 1000 µm.

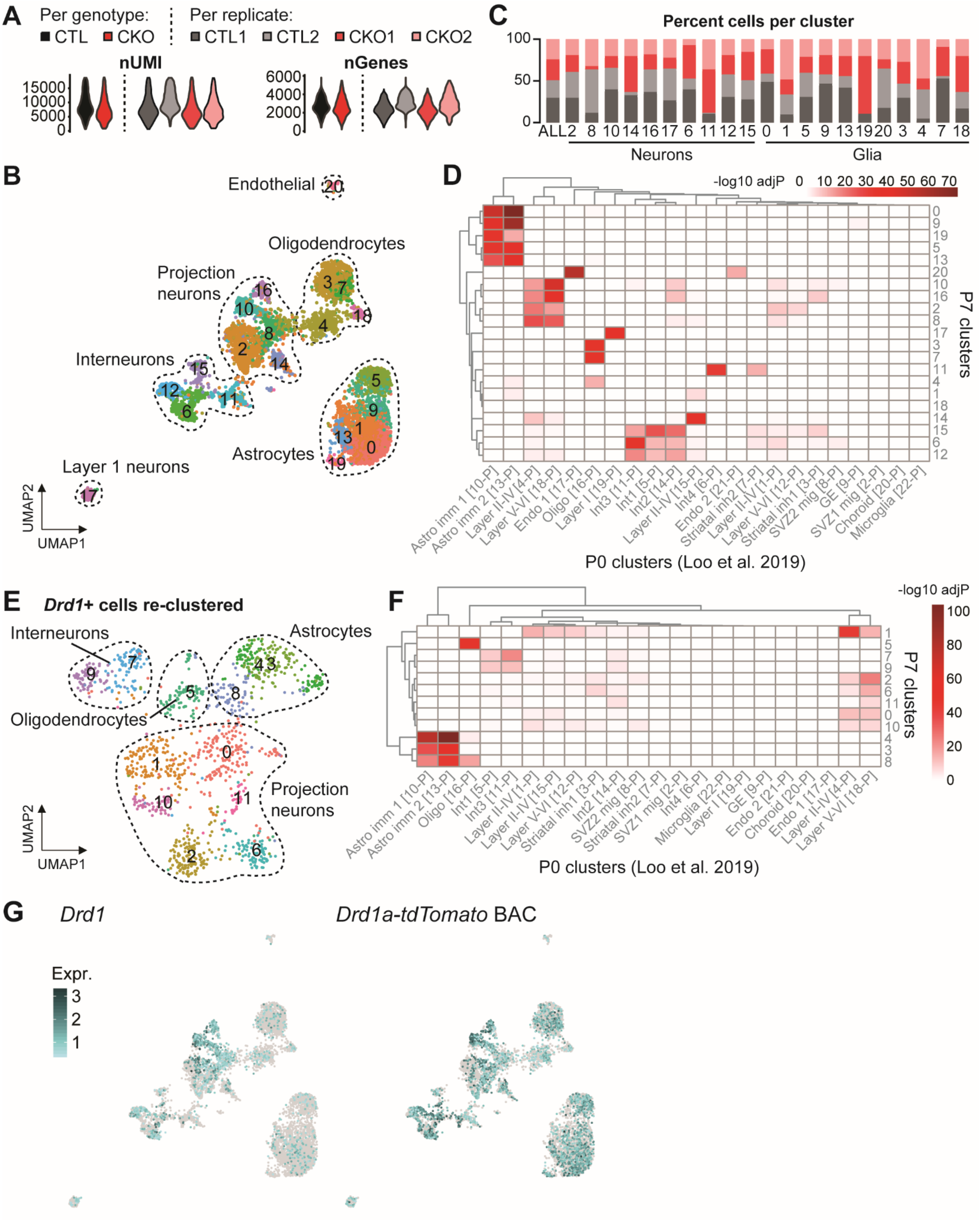
A
Per genotype:
CTL
CKO
Per replicate:
CTL1
CTL2
CKO1
CKO2
nUMI
nGenes
C
Percent cells per cluster
Neurons
Glia
B
Endothelial
Oligodendrocytes
Projection neurons
Interneurons
Astrocytes
Layer 1 neurons
UMAP2
UMAP1
D
-log10 adjP
P7 clusters
P0 clusters (Loo et al. 2019)
E
Drd1+ cells re-clustered
Interneurons
Astrocytes
Oligodendrocytes
Projection neurons
F
-log10 adjP
P7 clusters
P0 clusters (Loo et al. 2019)
G
Drd1
Drd1a-tdTomato BAC
Expr.

**Supplemental Figure 4. Single-cell RNA-seq analysis of FACS-isolated cells from postnatal *Drd1a*-tdTomato+ control and *Foxp2* cKO frontal cortex.** (A) Violin plots showing number of UMIs (left) and number of genes (right) detected per genotype and per replicate. (B) UMAP projection of clusters identified from control and cKO cells combined. (C) Percentage of cells from each replicate per cluster, colored based on (A). (D) Hypergeometric overlaps of marker genes from our P7 *Drd1a*-tdTomato clusters with P0 mouse cortex clusters from (Loo et al. 2019). (E) UMAP projection of re-clustered *Drd1*+ cells from (B). (F) Hypergeometric overlaps of *Drd1*+ cell cluster marker genes with P0 mouse cortex clusters from (Loo et al. 2019). (G) Feature plots of clusters from (B) showing expression levels of *Drd1* (left) and a partial *Drd1a*-tdTomato BAC sequence (right).

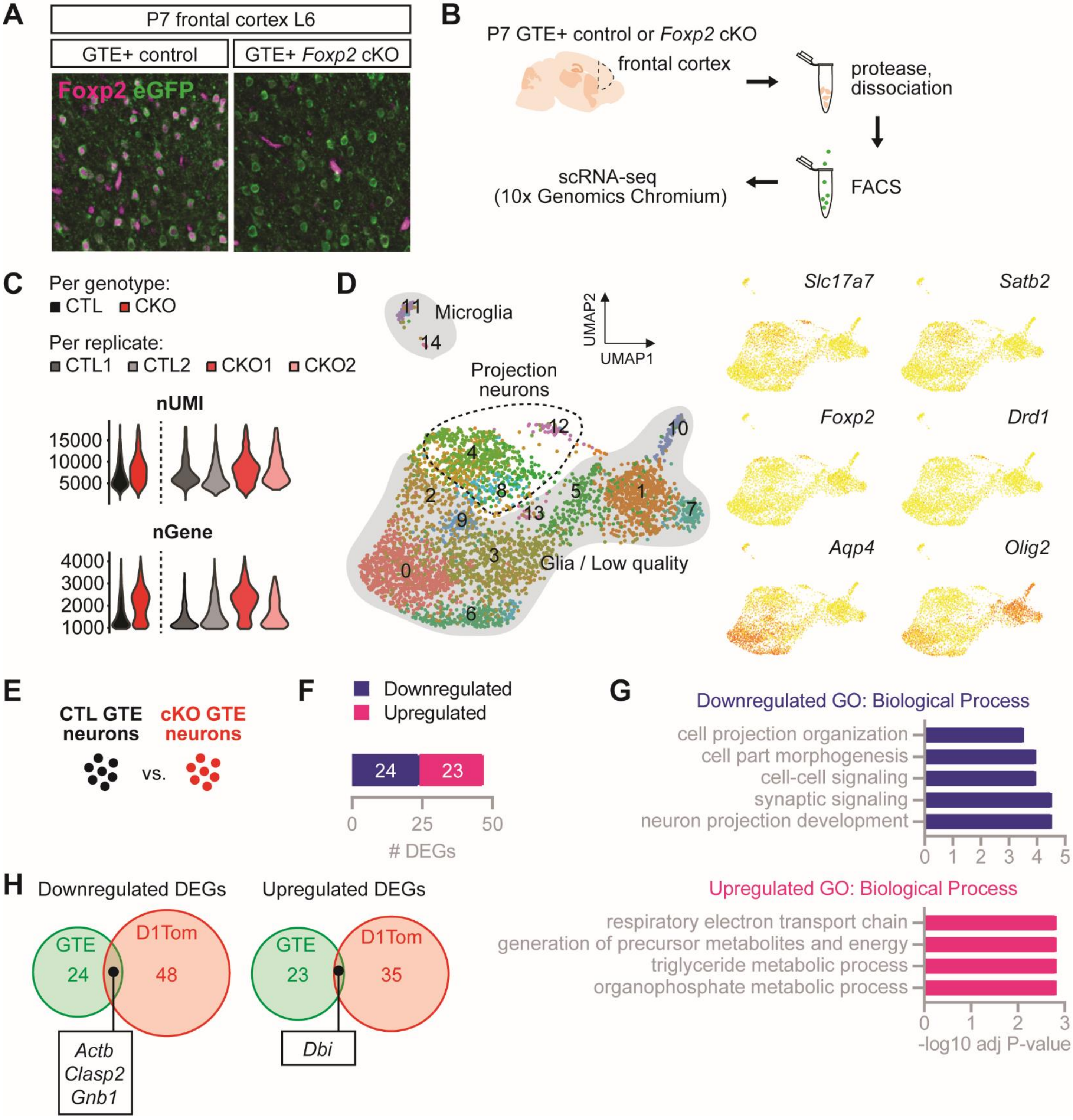


**Supplemental Figure 5. Single-cell RNA-seq analysis of FACS-isolated cells from postnatal *golli*-τ-eGFP+ control and *Foxp2* cKO frontal cortex.** (A) IHC for Foxp2 and eGFP in P7 *golli*-τ-eGFP+ (GTE+) control and *Foxp2* cKO frontal cortical L6. Residual Foxp2 signal in cKO cortex is non-specific blood vessel staining. (B) Experimental design for GTE scRNA-seq. *n* = 2 per condition. (C) Violin plots showing number of UMIs (top) and number of genes (bottom) detected per genotype and per replicate. (D) UMAP projection of cell type clusters (left) and marker gene feature plots (right) from control and cKO GTE scRNA-seq. (E) Identification of differentially expressed genes (DEGs) between all control and all *Foxp2* cKO neurons from GTE scRNA-seq. (F) Number of DEGs significantly down- or upregulated in cKO neurons. (G) Summarized gene ontology (GO) Biological Process terms for down- and upregulated DEGs. (H) Overlap of pseudo-bulk RNA-seq DEGs from the GTE and *Drd1a*-tdTomato (D1Tom) (Fig. 4B) analyses.

### Supplemental Methods

#### *Foxp2* cKO confirmation

For confirmation by IHC, sagittal sections of 20 µm thickness from male mice aged 14 weeks were stained using rabbit α-Foxp2 (#5337S, Cell Signaling Technology, 1:250) as described in the main Materials and Methods Immunohistochemistry section. Images were acquired using a Zeiss Axioscan Z1 slide scanner at the UT Southwestern Whole Brain Microscopy Facility and processed using Zeiss ZEN Lite and FIJI. Images were annotated using the Allen Mouse Brain Atlas. Confirmation by Western blotting was performed as described in the main Materials and Methods Western blotting section using the following antibodies: rabbit α-Foxp2 (#5337S, Cell Signaling Technology, 1:1000), mouse α-GAPDH (#MAB374, Millipore Sigma, 1:10,000), donkey α-rabbit IgG IRDye 800 (#926-32213, LI-COR Biosciences, 1:20,000), donkey α-mouse IgG IRDye 680 (#926-68072, LI-COR Biosciences, 1:20,000).

#### Behavioral analyses

Behavioral studies were performed in the following cohorts in the following order: Cohort 1 (neonatal USVs and motor tests, adult USVs), Cohort 2 (novel cage activity, open field, elevated plus maze), Cohort 3 (water Y-maze), Cohort 4 (spontaneous alternation). Mice in Cohorts 2-4 were habituated to handling for 5 min per day for 5 days the week prior to testing. All behavioral experiments were performed during the light cycle in the afternoon by an experimenter blinded to genotype.

#### Spontaneous alternation in T-maze

Mice were tested according to (Deacon and Rawlins 2006). After habituation to a dimly lit testing room for 15 min, the test mouse was placed in the start arm of the T-maze and allowed to enter a goal arm, where it was confined by a guillotine door for 30 s. The mouse was transferred to a holding cage for 15 s while the maze was wiped clean with diluted Process NPD (Steris Life Sciences) to remove odor cues. The mouse was then placed back in the start arm of the maze and allowed to choose a goal arm.

Failure to choose a goal arm within 2 min resulted in a failed trial and confinement to the previously chosen arm for 30 s. Mice were tested for 11 consecutive trials. Spontaneous alternation rate was calculated as number of alternations divided by number of opportunities to alternate (e.g. 10 if no failed trials) and multiplied by 100. Latency to goal arm was also measured. Comparisons of alternation rates between genotypes, and between each genotype and chance levels (50%), were performed using unpaired t-tests. Differences in latency to goal arm between genotypes were assessed using a two-way ANOVA with Bonferroni's multiple comparisons test.

**Novel cage activity**

Mice were tested according to (Araujo et al. 2017). Mice were placed into clean, plastic 18×28 cm cages with minimal bedding, and then each cage was then placed into a dark Plexiglas box. Photobeam Activity System-Home Cage software (San Diego Instruments) was used to record the number of infrared beam breaks for 2 h. Data were binned into 5 min intervals and differences between genotypes were assessed using a two-way ANOVA with Bonferroni's multiple comparisons test.

**Open field**

Mice were tested according to (Araujo et al. 2015). Mice were placed in a 16×16 in Plexiglass box and allowed to explore the arena for 10 min. Ethovision XT software (Noldus) was used to calculate the total distance moved and time spent in each zone of the field. Differences between genotypes were assessed using a two-tailed t-test.

**Elevated plus maze**

Mice were tested according to (Walf and Frye 2007). Mice were placed in the center of a plus-shaped maze with two open arms and two enclosed arms and video recorded for 5 min. Ethovision XT software (Noldus) was used to calculate the total distance moved and time spent in each zone of the maze. Differences between genotypes were assessed using a two-tailed t-test.

**Righting reflex**

Mice were tested according to (Araujo et al. 2015). Each pup was placed in a supine position on a clean, unobstructed surface, and the time taken to right onto all fours was measured. If a pup failed to right after 60 s, the time was recorded as 60 s. Each pup received one trial at each postnatal time point. Differences between genotypes were assessed using a two-way ANOVA with Bonferroni's multiple comparisons test.

**Negative geotaxis**

Mice were tested according to (Usui et al. 2017b). Each pup was oriented facing downward at a 30-degree angle on a sloped rough surface, and the time taken to reorient to face upward was recorded. If a pup took longer than 180 s, the time was recorded as 180 s. Each pup received one trial at each postnatal time point. Differences between genotypes were assessed using a two-way ANOVA with Bonferroni's multiple comparisons test.

**Cortical layer thickness analysis**

Cortical layer thickness was analyzed as described in (Usui et al. 2017a). Coronal sections of 30 µm thickness from male P7 mice were stained as described in the main Materials and Methods Immunohistochemistry section. For CTIP2 staining, citrate antigen retrieval was performed. The following antibodies and dilutions were used: rat α-CTIP2 (#ab18465, Abcam, 1:500), rabbit α-CUX1/CDP (#sc13024, Santa Cruz Biotechnology, 1:500), rabbit α-TBR1 (#ab31940, Abcam, 1:500). Images of primary somatosensory cortex (S1) were acquired using a Zeiss LSM 710 confocal laser scanning microscope at the UT Southwestern Neuroscience Microscopy Facility and processed and analyzed using Zeiss ZEN Lite and FIJI. For each animal, cortical thickness was calculated as the average of 3 measurements of S1 thickness. Then each layer thickness was calculated as the average of 3 measurements based on cell architecture (DAPI-based) or layer marker staining (Marker-based). Relative layer thickness was calculated as layer thickness divided by cortical thickness multiplied by 100, and compared between genotypes using a two-tailed t-test.

**Cortical projection analysis**

To visualize cortical layer 6 axons, *golli-τ-eGFP* transgenic mice from (Jacobs et al. 2007) were backcrossed with C57BL/6J mice for at least 10 generations and mated with *Emx1-Cre* mice and *Foxp2*$^{flox/flox}$ mice to produce *Foxp2* cKO and control mice carrying the transgene. Coronal sections of 40 µm thickness from male and female P14 mice were stained as described in the main Materials and Methods Immunohistochemistry section using chicken α-GFP (#GFP-1010, Aves Labs, 1:1000). Images were acquired using a Zeiss Axioscan Z1 slide scanner at the UT Southwestern Whole Brain Microscopy Facility and processed using Zeiss ZEN Lite and FIJI.

**Single-cell RNA-seq**

***Drd1*+ cell clustering**

Cells with *Drd1* UMI >1 were pulled from the full *Drd1a*-tdTomato dataset and re-clustered with resolution 1.6, then annotated as described in the main Materials and Methods scRNA-seq section.

***Drd1a-tdTomato* BAC mapping**

A partial *Drd1a-tdTomato* BAC sequence containing the *tdTomato* coding sequence and residual cloning vector sequences was obtained from Dr. Bernd Gloss (Ade et al. 2011). *makeblastdb* (Magic-BLAST v1.4.0) was used to create an index for the BAC sequence fasta file (Boratyn et al. 2018). Extracted fastq files for selected/whitelisted cell barcodes were then aligned to the BAC sequence index using *magicblast* (Magic-BLAST v1.4.0). *featureCounts* (Subread v1.6.2) was used to count the aligned reads to the BAC sequence. After sorting the *featureCounts* resulting BAM file using Samtools v1.6, reads for the BAC sequence were counted across cells using *count* (UMI Tools v0.5.4) to generate a count table similar to a raw UMI table. The tdTomato counts table was merged with the existing gene count table, and the combined raw count tables were filtered for cells passing the following thresholds (to obtain the same cells as the original *Drd1a*-tdTomato scRNA-seq analysis): <20,000 UMIs, <10%

mitochondrial transcripts, <20% ribosomal protein gene transcripts. The counts were normalized using Seurat's *NormalizeData* function, and the normalized expression was used for the *Drd1a*-tdTomato BAC feature plot.

#### *Golli*-τ-eGFP (GTE) dataset

eGFP+ cells were isolated from P7 *golli*-τ-eGFP control and *Foxp2* cKO frontal cortex by FACS and used to prepare libraries as described in the main Materials and Methods scRNA-seq section. Each sample was prepared on separate days: GTE-CTL1 (M), GTE-CTL2 (M), GTE-CKO1 (M), GTE-CKO2 (F). Libraries were pooled and sequenced twice using an Illumina NextSeq 500 at the McDermott Sequencing Core at UT Southwestern. Clustering, annotation, and pseudo-bulk DEG calculation, and GO analyses were performed as described in the main Materials and Methods, with a clustering resolution of 1.4.
